# The anterior insular cortex mediates higher-order threat learning and memory

**DOI:** 10.64898/2026.10.09.757763

**Authors:** Ho Namkung, Bandhan Mukherjee, Josephine de Chabot, Sedona Lockhart, Lauren Guttman, Imad Isehak, Annatje Clark, Atsushi Saito, Isamu Miura, Koko Ishizuka, Hyung-Bae Kwon, Akira Sawa

## Abstract

Learning about potential threats is indispensable for survival, and its deficits underlie numerous neuropsychiatric disorders. Human brain imaging suggests the anterior insular cortex (AIC) as a key neural correlate of threat learning, yet its mechanistic role remains unclear. Leveraging multimodal systems neuroscience approaches, we demonstrate that AIC excitatory neurons, specifically layer 5 pyramidal neurons projecting to the basolateral amygdala, are essential for higher-order threat learning/memory. They are specifically required when a predictor is temporally separated from an aversive event or when the predictive information is embedded within a diffuse context. In contrast, the AIC is dispensable for elementary, temporally contiguous threat learning. Surprisingly, while the AIC promotes memory formation, it also constrains the level of fear expression during retrieval in a context-dependent manner. This bidirectional control is mediated by the recruitment of distinct AIC neuronal ensembles. Together, we identify a novel circuit mechanism for bidirectional regulation of higher-order threat learning/memory.

## INTRODUCTION

The ability to detect and learn about potential threats in the environment is indispensable for survival. This process, known as threat learning, relies on the brain’s capacity to identify and encode key predictors of danger, thereby guiding future defensive behaviors^1–3^. Deficits in threat learning are central to the pathophysiology of numerous neuropsychiatric disorders, underscoring the clinical significance of understanding its underlying mechanisms^4–9^. The neural substrates of threat learning have been extensively investigated using fear conditioning paradigms in both humans and animal models^8,10–14^. In these paradigms, a neutral stimulus (conditioned stimulus, CS), such as a discrete cue or a broader environmental context, acquires emotional salience through its association with an aversive event (unconditioned stimulus, US)^8,11^. This association transforms the CS into a potent predictor of threat, eliciting defensive responses even in the absence of the US^8,11^.

Fear conditioning (FC) paradigms are broadly categorized into cued and contextual FC, distinguished by the nature of the CS^8,15,16^. In cued FC, the CS is a discrete, temporally defined stimulus, such as a tone or light. Cued FC can be further subdivided into delay and trace FC, based on the presence of a temporal interval between the CS and US^8,17^. In delay FC, the US co-terminates with or immediately follows the CS. In contrast, trace FC introduces a time interval (TI) between the termination of the CS and the onset of the US, requiring the brain to maintain a “trace” of the CS during the interval to form the association. Due to this additional temporal demand, trace FC may recruit higher-order cognitive processes^18–21^.

Contextual FC involves the association of the US with a diffuse, multimodal context rather than a discrete cue^8,13^. This requires the integration of distributed sensory features into a cohesive environmental representation, and previous studies have indicated the involvement of higher-order neural processing associated with the cortical and hippocampal regions^13,22^. By systematically applying delay, trace, and contextual FC paradigms, researchers can compare the neural circuit mechanisms underlying both elementary and complex forms of threat learning.

Human fMRI studies have identified key brain regions consistently activated during cued and/or contextual FC, including the amygdala, anterior cingulate cortex (ACC), hippocampus, and insular cortex^23–32^. While the roles of the amygdala, ACC, and hippocampus in fear learning and memory have been extensively characterized, the precise function of the insular cortex remains poorly understood. This gap in knowledge is likely due to the relatively limited number of animal studies focusing on the insular cortex, which are essential for addressing causal and mechanistic questions^33^. Fortunately, a recent surge of interest in the insular cortex in animal models, particularly mice, has begun to address this gap^34^.

However, these efforts have primarily focused on characterizing the posterior insular cortex (PIC) in fear learning and memory^35–44^, whereas studies of the anterior insular cortex (AIC) remain relatively scarce. Notably, the AIC is not cytoarchitectonically uniform along its rostrocaudal axis: its caudal portion contains agranular, dysgranular, and granular subdivisions. In contrast, its rostral portion consists exclusively of agranular cortex and lacks layer IV. Existing AIC studies of fear learning have largely targeted the caudal AIC and have studied the elementary threat learning using the delay FC paradigm^45–47^. Consequently, the contribution of the rostral AIC, a purely agranular region that is distinct from caudal AIC in its cytoarchitecture and connectivity^33,34,48^, remains largely unexplored. This gap remains particularly in more complex forms of fear learning and memory, such as trace and contextual FC.

Here, we address this gap by systematically characterizing the role of the AIC, particularly its rostral agranular subregion, across three distinct FC paradigms (delay, trace, and contextual) and their respective memory phases (acquisition, consolidation, and retrieval). We first compare the AIC’s involvement in delay and trace FC paradigms, which differ by the presence of a TI between the CS and US. Building on the comparative insights from these two paradigms, we then extend our investigation to the contextual FC paradigm. By using multimodal techniques, we investigated the causal role of AIC in a paradigm-, memory phase-, cell type-, and neural circuit-specific manner.

## RESULTS

Schematic overview of FC paradigms and corresponding memory phases are summarized in **Figure S1**.

### The AIC is required for fear memory acquisition in trace FC, but not in delay FC

We first questioned whether AIC activity is necessary for fear memory acquisition in delay and trace FC. To address this question, we bilaterally infused a mixture of muscimol (GABA_A r_eceptor agonist) and baclofen (GABA_B r_eceptor agonist) into the AIC 30 min before the conditioning session to transiently inactivate this region (**Figures 1A and S2A**)^49^. In delay FC, AIC inactivation had no effect on fear responses to the CS during the retrieval session 24 h later (**Figures 1A and 1B**). In trace FC, in contrast, AIC inactivation significantly reduced fear responses to the CS during the retrieval session (**Figures 1A and 1C**). These findings suggest that AIC activity is required for fear memory acquisition in trace FC, but not in delay FC.

**Figure 1.**
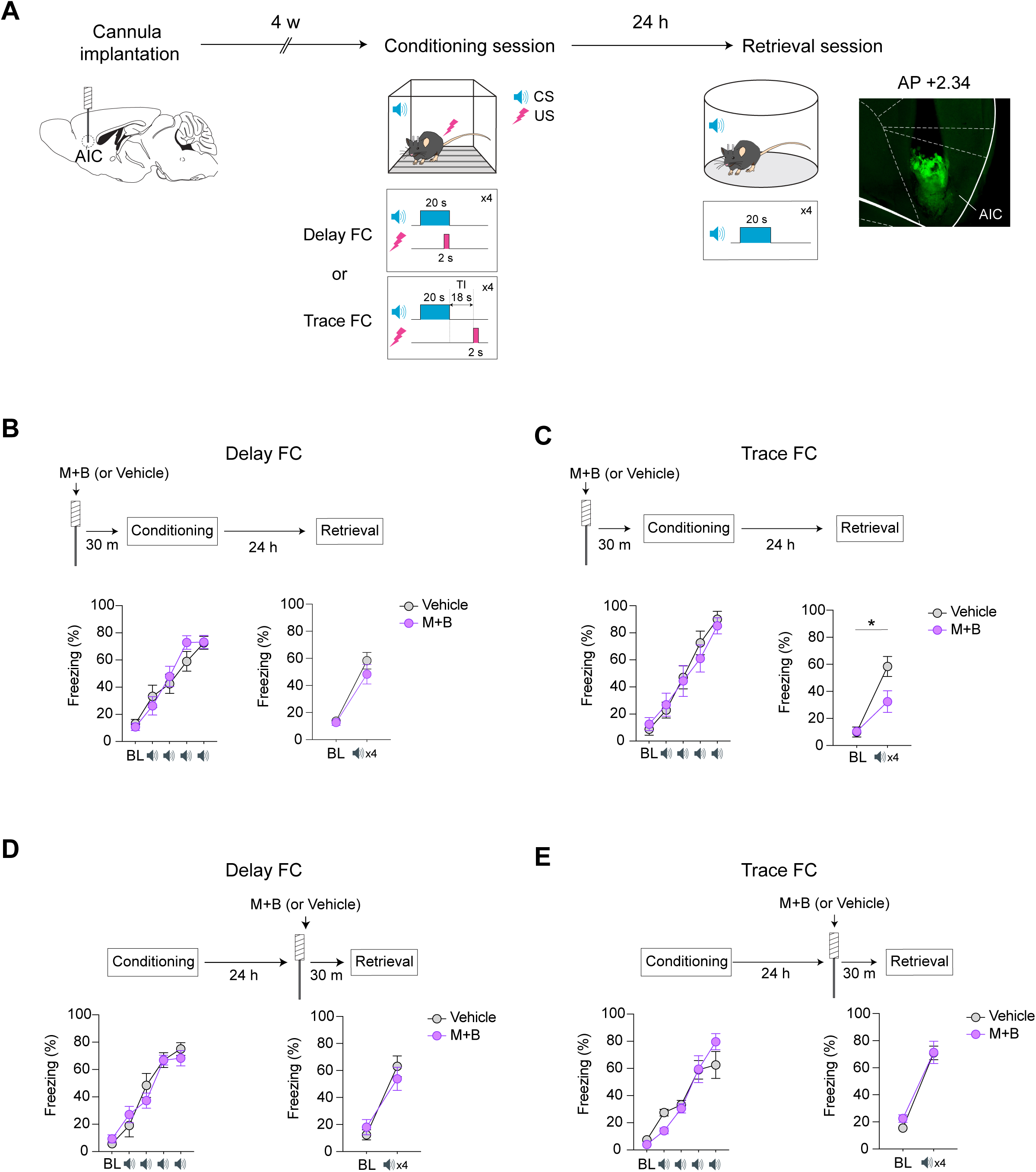
AIC activity is required for fear memory acquisition in trace FC, but not in delay FC. **(A)** Left: Schematic of pharmacological inhibition experiments conducted in cued FC (delay FC and trace FC). Right: Representative histological image showing the approximate location of the cannula tip and the extent of drug diffusion. **(B)** In delay FC, pharmacological inactivation of the AIC did not affect fear responses to the CS during the retrieval session. Baseline (BL) represents the mean freezing during the initial 2-minute period prior to the first CS. Retrieval performance was quantified as the average freezing across four CS presentations in a neutral context. Conditioning (*N*_Vehicle=_12, *N*_M+B=_12; two-way mixed ANOVA, *p*=0.235). Retrieval (*N*_Vehicle=_12, *N*_M+B=_12; two-way mixed ANOVA, *p*=0.411). **(C)** In trace FC, pharmacological inactivation of the AIC significantly reduced fear responses during the retrieval session. Conditioning (*N*_Vehicle=_11, *N*_M+B=_13; two-way mixed ANOVA, *p*=0.617). Retrieval (*N*_Vehicle=_11, *N*_M+B=_13; two-way mixed ANOVA, *p*=0.01). **(D)** In delay FC, pharmacological inactivation of the AIC during the retrieval session did not affect fear responses. Conditioning (*N*_Vehicle=_10, *N*_M+B=_12; two-way mixed ANOVA, *p*=0.360). Retrieval (*N*_Vehicle=_10, *N*_M+B=_12; two-way mixed ANOVA, *p*=0.214). **(E)** In trace FC, pharmacological inactivation of the AIC during the retrieval session had no impact on fear responses. Conditioning (*N*_Vehicle=_10, *N*_M+B=_10; two-way mixed ANOVA, *p*=0.130). Retrieval (*N*_Vehicle=_10, *N*_M+B=_10; two-way mixed ANOVA, *p*=0.559). \**p* < 0.05. Graph expressed as mean ± SEM. More detailed statistical information is in Table S1.

Since pre-conditioning inactivation may have a lasting impact on AIC activity during the post-conditioning period^50^, we next investigated whether the observed impairment was due to a disruption of AIC-mediated memory consolidation. To test this, a mixture of muscimol and baclofen was infused bilaterally into the AIC immediately after the conditioning session in trace FC. This post-conditioning inactivation did not affect fear responses during the retrieval session (**Figure S2B**), consistent with the result from post-conditioning chemogenetic inhibition of AIC excitatory neurons using the inhibitory designer receptors exclusively activated by designer drugs (DREADD) hM4Di (**Figure S2C**)^51^. These findings indicate that AIC activity is required for memory acquisition, but not for memory consolidation, processes in trace FC.

Finally, we assessed whether AIC activity is necessary for fear memory retrieval by infusing muscimol and baclofen 30 min before the retrieval session. This manipulation did not affect fear responses to the CS during retrieval in either delay or trace FC (**Figures 1D and 1E**). In summary, AIC activity is required for fear memory acquisition in trace FC, but not in delay FC, with its involvement not extending to memory consolidation or retrieval.

### Activity of AIC excitatory neurons during the trace interval (TI) is required for fear memory acquisition in trace FC

Although pharmacological inactivation with muscimol and baclofen, affecting both excitatory and inhibitory neurons^49^, is useful for an initial screening, it does not offer cell type-specificity and high temporal precision. To overcome these limitations and further refine our mechanistic understanding, we employed temporally specific optogenetic manipulation to selectively inhibit excitatory neurons within the AIC^52^. We focused on excitatory neurons given their fundamental role in orchestrating long-range communication and mediating the synaptic remodeling within the distributed neural networks that subserve fear memory^48^. We used enhanced halorhodopsin (eNpHR3.0) expressed under the CaMKIIα promoter to optogenetically inhibit AIC excitatory neurons during specific time windows in trace FC: the CS, TI, US, or the entire trial period encompassing the CS-TI-US sequence (**Figure 2A**).

**Figure 2.**
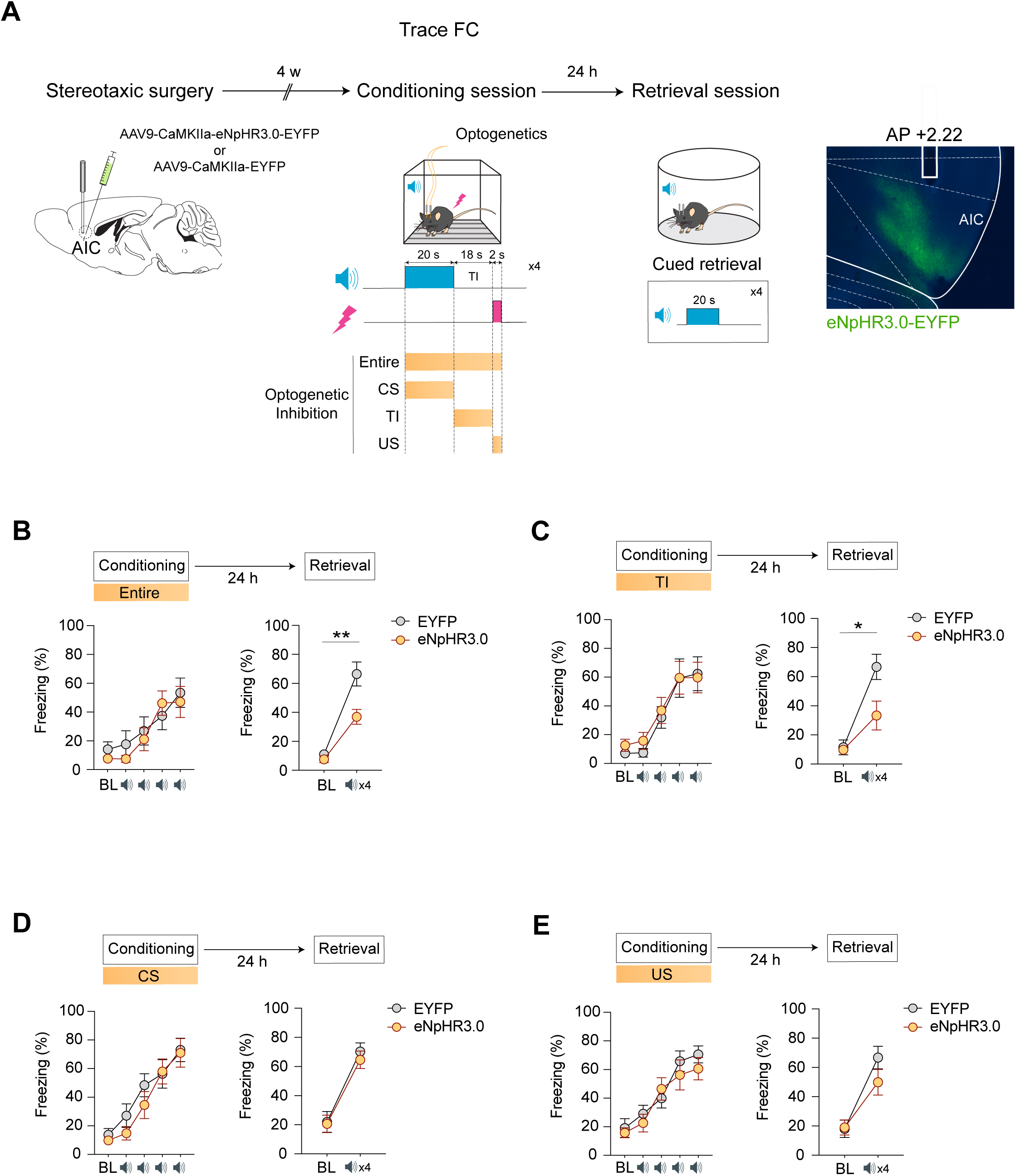
Activity of AIC excitatory neurons specifically during the trace interval (TI) is required for fear memory acquisition in trace FC. **(A)** Left: Schematic of temporally specific optogenetic inhibition experiments. Right: Representative histological image showing the approximate cannula tip location and the extent of AAV-mediated eNpHR3.0-EYFP expression in AIC excitatory neurons. **(B)** Optogenetic inhibition of AIC excitatory neurons throughout the entire CS-TI-US period prevented fear memory acquisition. Conditioning (*N*_EYFP=_10, *N*_eNpHR3.0=_10; two-way mixed ANOVA, *p*=0.591). Retrieval (*N*_EYFP=_10, *N*_eNpHR3.0=_10; two-way mixed ANOVA, *p*=0.009). **(C)** Optogenetic inhibition of AIC excitatory neurons during the TI impaired fear memory acquisition. Conditioning (*N*_EYFP=_10, *N*_eNpHR3.0=_10; two-way mixed ANOVA, *p*=0.868). Retrieval (*N*_EYFP=_10, *N*_eNpHR3.0=_10; two-way mixed ANOVA, *p*=0.026). **(D)** Optogenetic inhibition of AIC excitatory neurons during the CS did not affect fear memory acquisition. Conditioning (*N*_EYFP=_11, *N*_eNpHR3.0=_11; two-way mixed ANOVA, *p*=0.700). Retrieval (*N*_EYFP=_11, *N*_eNpHR3.0=_11; two-way mixed ANOVA, *p*=0.685). **(E)** Optogenetic inhibition of AIC excitatory neurons during the US did not significantly impact fear memory acquisition. Conditioning (*N*_EYFP=_11, *N*_eNpHR3.0=_11; two-way mixed ANOVA, *p*=0.470). Retrieval (*N*_EYFP=_11, *N*_eNpHR3.0=_11; two-way mixed ANOVA, *p*=0.132). \**p* < 0.05, \*\**p < 0.01. Data are expressed as mean ± SEM.* More detailed statistical information is in Table S1.

Inhibition throughout the entire trial period prevented fear memory acquisition, consistent with the effect of pharmacological AIC inactivation (**Figure 2B**). Notably, selective inhibition during the TI alone was sufficient to prevent fear memory acquisition (**Figure 2C**), whereas inhibition during the CS or US had no significant effect (**Figures 2D and 2E**). Together, these results demonstrate that AIC excitatory neuron activity specifically during the TI is required for fear memory acquisition. In contrast, optogenetic inhibition of PIC excitatory neurons did not disrupt fear memory acquisition (**Figures S3A and S3B**), highlighting the distinct role of the AIC compared to the PIC.

Given the critical role of AIC excitatory neurons during the TI, we next examined their activity dynamics throughout the course of trace FC, with a particular focus on learning-related changes during the TI. We employed *in vivo* fiber photometry to record population-level activity of AIC excitatory neurons, using the fluorescent calcium indicator GCaMP6f expressed under the CaMKIIα promoter (**Figures 3A and 3B**). Intriguingly, across four CS-US pairings (Trial 1 -Trial 4) during the conditioning session, AIC excitatory neurons developed an increasingly prominent transient response at the onset of the TI (**Figure 3C**). Accordingly, their mean activity during the TI significantly increased across trials, whereas activity during the CS and US did not significantly change (**Figure 3D**). Although AIC excitatory neurons were also activated during the CS and US, our previous optogenetic data demonstrate that these specific activities do not play a causal role in fear memory acquisition (**Figures 2C and 2E**). Collectively, the emergence of activity in AIC excitatory neurons specifically during the TI likely represents a neural correlate of successful fear memory acquisition.

**Figure 3.**
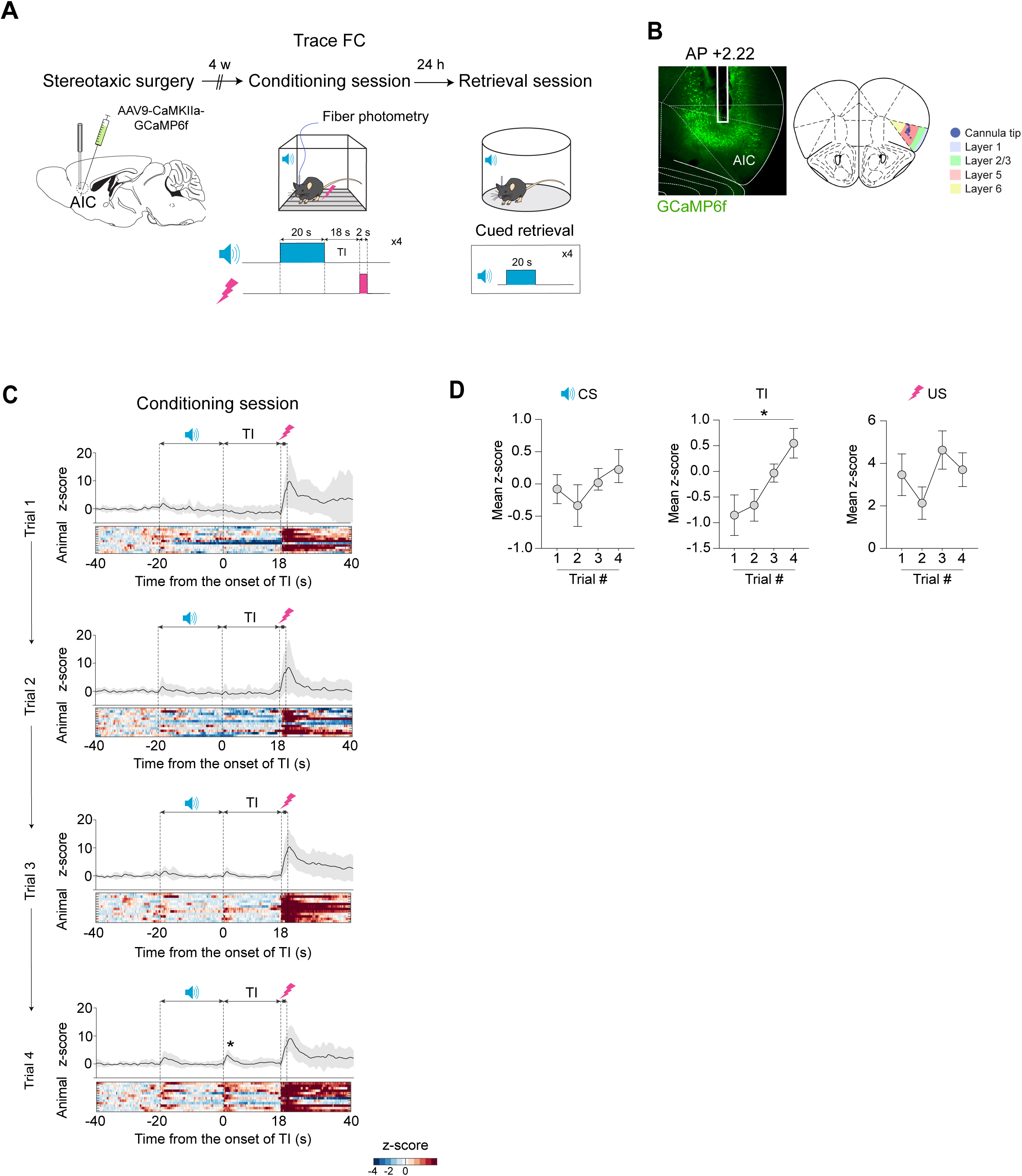
AIC excitatory neurons encode a learning-related activity signature at the onset of the TI that becomes increasingly prominent across trials. **(A)** Schematic of the fiber photometry experiments in trace FC. **(B)** Left: Representative histological image showing the approximate location of the cannula tip and the extent of AAV-mediated GCaMP6f expression in AIC excitatory neurons. Right: Histological assessment of the fiber optic cannula implantation sites indicates primarily layer 5 targeting. **(C)** Population-level recordings of AIC excitatory neurons reveal the emergence of a transient activity (indicated by an asterisk), time-locked to the onset of the TI, across four CS-US pairing trials (Trial 1 -Trial 4). **(D)** Across the trials, mean activity significantly increased during the TI, but not during the CS or US. CS (*N*=12; one-way repeated measures ANOVA, *p*=0.552). TI (*N*=12; one-way repeated measures ANOVA, *p*=0.016). US (*N*=12; one-way repeated measures ANOVA, *p*=0.179). \**p* < 0.05, *Data are expressed as mean ± SEM.* More detailed statistical information is in Table S1.

### Activity of AIC L5 pyramidal neurons (L5-PNs) is required for fear memory acquisition in trace FC

We next addressed whether AIC excitatory neurons engaged during the TI may be layer-specific. We employed soma-targeted Cal-Light (ST-Cal-Light)^53^, a recently developed, temporally specific, activity-dependent labeling technique, which enables the selective expression of the reporter enhanced green fluorescent protein (EGFP) in AIC excitatory neurons that are specifically active during the TI under blue-light control (**Figure 4A**). Using this approach, we found that excitatory neurons active during the TI are predominantly enriched in the area that corresponds to the layer 5 (L5) (**Figure 4B**).

**Figure 4.**
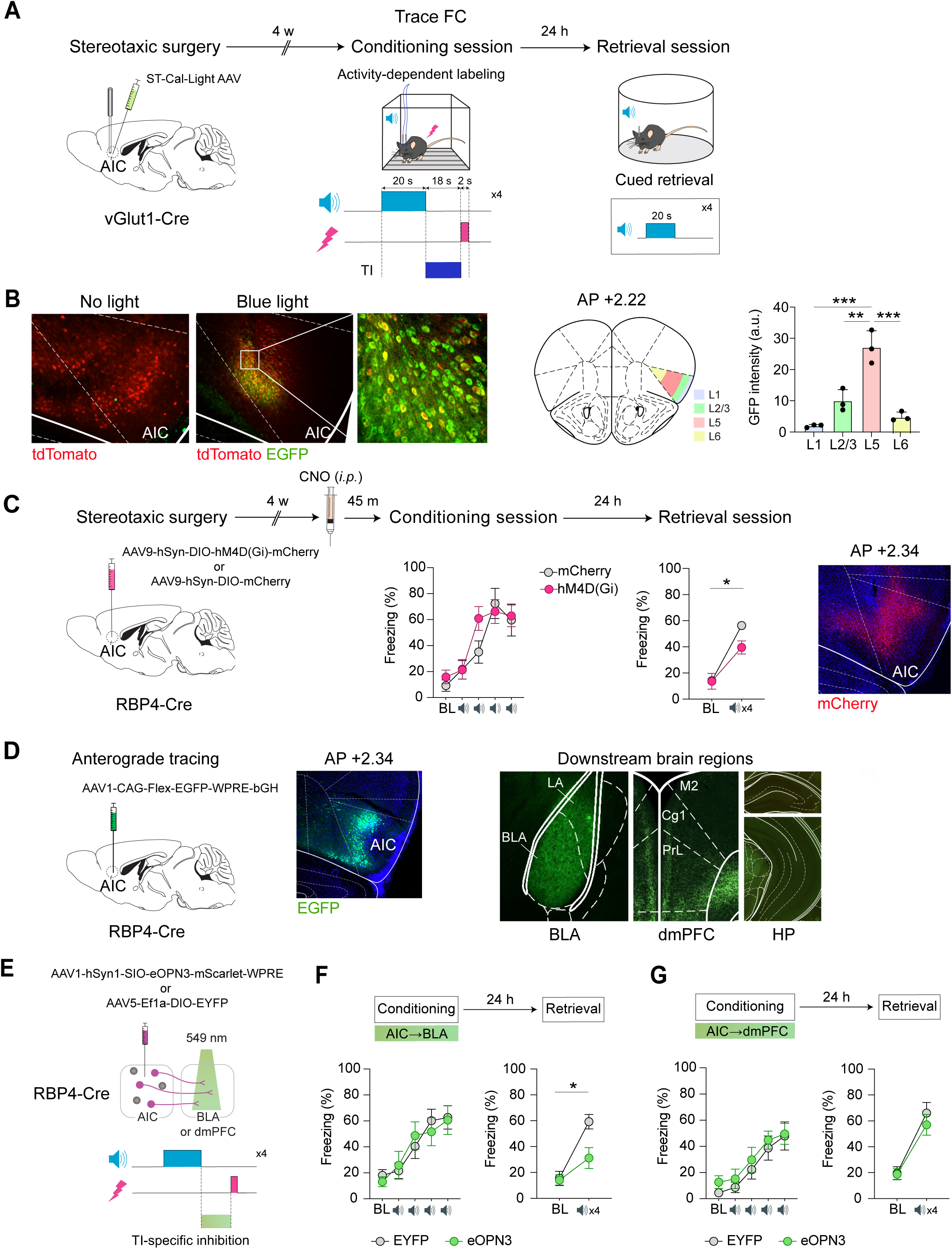
TI-specific activity of AIC L5-PNs, projecting to the basolateral amygdala, is critical for fear memory acquisition in trace FC. **(A)** Schematic of the temporally specific ST-Cal-Light labeling experiments, designed to tag AIC excitatory neurons active during the TI. **(B)** Left: Two representative histological images. In the first image, ST-Cal-Light AAV-injected cells (red) were shown without blue-light activation, thus remaining unlabeled. In the second image, excitatory neurons active specifically during the TI (green) were labeled under blue-light illumination. Right: AIC excitatory neurons active during the TI were predominantly enriched in layer 5 (L5). Layers (*N*=3; one-way ANOVA, *p*=7.176×10^-5^; post-hoc Tukey’s HSD). **(C)** Chemogenetic inhibition of AIC L5-PNs during the conditioning session disrupted fear memory acquisition, consistent with the effects from optogenetic and pharmacological inhibition. Conditioning (*N*_mCherry=_8, *N*_hM4D(Gi)=_8; two-way mixed ANOVA, *p*=0.220). Retrieval (*N*_mCherry=_8, *N*_hM4D(Gi)=_8; two-way mixed ANOVA, *p*=0.037). **(D)** Left: Anterograde tracing of AIC L5-PNs. Right: AIC L5-PNs project robustly to the basolateral amygdala (BLA) and dorsomedial prefrontal cortex (dmPFC), but not to the hippocampus (HP). **(E)** Schematic of projection-specific optogenetic inhibition experiments. **(F)** Optogenetic inhibition of the AIC^L5-PNs^→BLA pathway during the TI impaired fear memory acquisition. Conditioning (*N*_EYFP=_10, *N*_eOPN3=_10; two-way mixed ANOVA, *p*=0.704). Retrieval (*N*_EYFP=_10, *N*_eOPN3=_10; two-way mixed ANOVA, *p*=0.025). **(G)** Optogenetic inhibition of the AIC ^L5-PNs^→dmPFC pathway during the TI had no effect on fear memory acquisition. Conditioning (*N*_EYFP=_12, *N*_eOPN3=_12; two-way mixed ANOVA, *p*=0.937). Retrieval (*N*_EYFP=_12, *N*_eOPN3=_12; two-way mixed ANOVA, *p*=0.456). \**p* < 0.05, \*\**p < 0.01,* \*\*\**p < 0.001. Data are expressed as mean ± SEM.* More detailed statistical information is in Table S1.

To further validate this finding *in vivo*, we performed fiber photometry to directly record the activity of L5 pyramidal neurons (L5-PNs) in the AIC. For this experiment, we expressed Cre-dependent GCaMP6f in the AIC of Retinol Binding Protein 4 (RBP4)-Cre mice^54^, a well-established line for selectively targeting cortical L5-PNs (**Figure S4A**). Consistent with our broader excitatory population recordings, AIC L5-PNs developed an increasingly prominent transient response at the onset of the TI across trials (Trial 1 -Trial 4), accompanied by a significant increase in their mean activity during the TI, but not during the CS or US, across trials (**Figures S4B and S4C**). These consistent findings likely arise because most fiber optic cannulas in our earlier experiments were positioned within L5, as confirmed by histological analysis (**Figure 3B**). Collectively, these findings suggest that AIC L5-PNs play a pivotal role in the emergence of activity time-locked to the onset of the TI.

We next tested whether selectively inhibiting AIC L5-PNs could recapitulate the behavioral effects of our earlier optogenetic and pharmacological inhibition. Indeed, using the inhibitory DREADD hM4Di, we found that chemogenetic inhibition of AIC L5-PNs during the conditioning session significantly reduced fear responses during the retrieval session (**Figure 4C**). Taken together, these data identify AIC L5-PNs as a key neuronal population that plays a causal role in fear memory acquisition in trace FC.

### TI-specific activity of AIC L5-PNs, projecting to the basolateral amygdala, is required for fear memory acquisition in trace FC

Next, we aimed to elucidate the neural circuit mechanisms by which AIC L5-PNs contribute to fear memory acquisition in trace FC. We hypothesized that AIC L5-PN activity during the TI engages specific downstream regions to cooperatively drive fear memory acquisition. Based on previous studies^8,18,48,55^, we identified three candidate regions as potential downstream targets: the basolateral amygdala (BLA), dorsomedial prefrontal cortex (dmPFC), and hippocampus (HP). Anterograde tracing revealed robust axonal projections from AIC L5-PNs to the BLA and dmPFC but not the HP, allowing us to rule out the HP from further investigation as a direct projection target (**Figure 4D**).

Consequently, we examined whether excitatory inputs from AIC L5-PNs to either the BLA or dmPFC are functionally required for fear memory acquisition in trace FC. To test this, we employed projection-specific optogenetic inhibition by expressing Cre-dependent inhibitory opsin eOPN3 in the AIC of RBP4-Cre mice and selectively silencing axonal terminals in either the BLA or dmPFC during the TI (**Figure 4E**). Notably, inhibition of the AIC^L5-PNs^→BLA pathway during the TI impaired fear memory acquisition (**Figure 4F**), whereas inhibition of the AIC ^L5-PNs^→dmPFC pathway had no effect (**Figure 4G**). These findings indicate that the activity of AIC L5-PNs projecting to the BLA, specifically during the TI, is critical for fear memory acquisition in trace FC.

### Dual roles of AIC excitatory neurons in contextual FC

While the AIC is not involved in delay FC, it plays a pivotal role in trace FC where a TI introduces temporal uncertainty about the timing of the US. Meanwhile, during contextual FC, the US occurs unpredictably within a diffuse environment, without a discrete cue to signal its timing. Accordingly, we hypothesized that the AIC may also play a crucial role in memory acquisition in contextual FC.

To address this question, we bilaterally infused a mixture of muscimol and baclofen into the AIC 30 min before the conditioning session in contextual FC. This manipulation significantly reduced fear responses to the conditioning context during the retrieval session (**Figures 5A and 5B**). We next examined whether this impairment resulted from a disruption of AIC-mediated consolidation processes that may have been persistently affected by pre-conditioning inactivation. To address this possibility, we expressed the inhibitory DREADD hM4Di in AIC excitatory neurons under the CaMKIIα promoter and chemogenetically inhibited these neurons immediately after the conditioning session (**Figure S2D**). Unlike pre-conditioning pharmacological inactivation, this post-conditioning manipulation had no effect on fear responses during the retrieval session. These results suggest that AIC activity is required for fear memory acquisition, but not for consolidation, in contextual FC.

**Figure 5.**
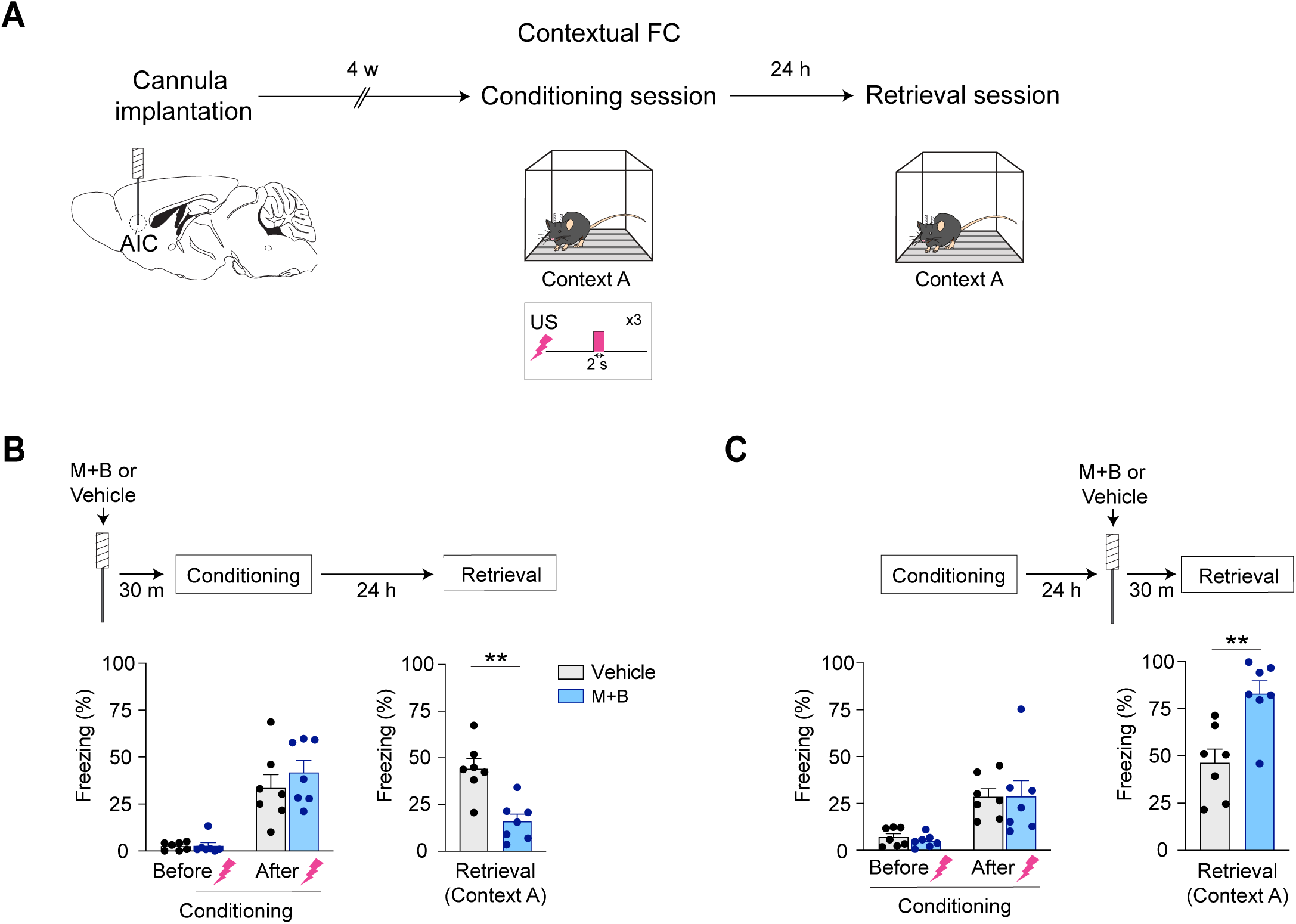
AIC activity is required for contextual fear memory acquisition and unexpectedly constrains fear responses during retrieval. **(A)** Schematic of pharmacological inhibition experiments conducted in contextual FC. **(B)** In contextual FC, pharmacological inactivation of the AIC during the conditioning session significantly reduced fear responses during the retrieval session. Conditioning (*N*_Vehicle=_7, *N*_M+B=_7; two-way mixed ANOVA, *p*=0.454). Retrieval (*N*_Vehicle=_7, *N*_M+B=_7; two-tailed Student’s *t*-test, *p*=0.001). **(C)** Pharmacological inactivation of the AIC during the retrieval session unexpectedly increased fear responses. Conditioning (*N*_Vehicle=_7, *N*_M+B=_7; two-way mixed ANOVA, *p*=0.784). Retrieval (*N*_Vehicle=_7, *N*_M+B=_7; two-tailed Student’s *t*-test, *p*=0.003). \*\**p < 0.01. Data are expressed as mean ± SEM.* More detailed statistical information is in Table S1.

Finally, we tested whether AIC activity is necessary for fear memory retrieval by infusing muscimol and baclofen 30 minutes before the retrieval session. Surprisingly, this treatment led to a significant increase in fear responses during the retrieval session, indicating that AIC activity may normally suppress fear responses during memory retrieval in contextual FC (**Figures 5A and 5C**). Altogether, these findings demonstrate that the AIC is involved in fear memory acquisition in both trace and contextual FC as hypothesized, while also revealing an unexpected role for the AIC in memory retrieval in contextual FC.

### AIC L5-PNs, projecting to the BLA, are required for fear memory acquisition in contextual FC

We next assessed the activity dynamics of AIC excitatory neurons during the conditioning session in contextual FC. We performed *in vivo* fiber photometry to record population-level activity using GCaMP6f expressed under the CaMKIIα promoter (**Figure 6A**). During the conditioning session, AIC excitatory neurons exhibited robust, time-locked responses to US delivery across all three context-US pairings (**Figure 6B**). To determine whether this activity is causally required for contextual fear memory acquisition, we used a temporally precise, cell type-specific optogenetic approach. We inhibited AIC excitatory neurons specifically around each context-US pairing (6-second windows covering each 2-second US) by photo-stimulating eNpHR3.0 expressed under the CaMKIIα promoter (**Figure 6C**). This manipulation significantly reduced fear responses to the conditioning context (Context A; **Figure 6D**), but not to the neutral context (Context B; **Figures S5A and S5B**), during the retrieval session, consistent with the pharmacological effect. These findings indicate that AIC excitatory neuron activity during context-US pairings is required for contextual fear memory acquisition.

**Figure 6.**
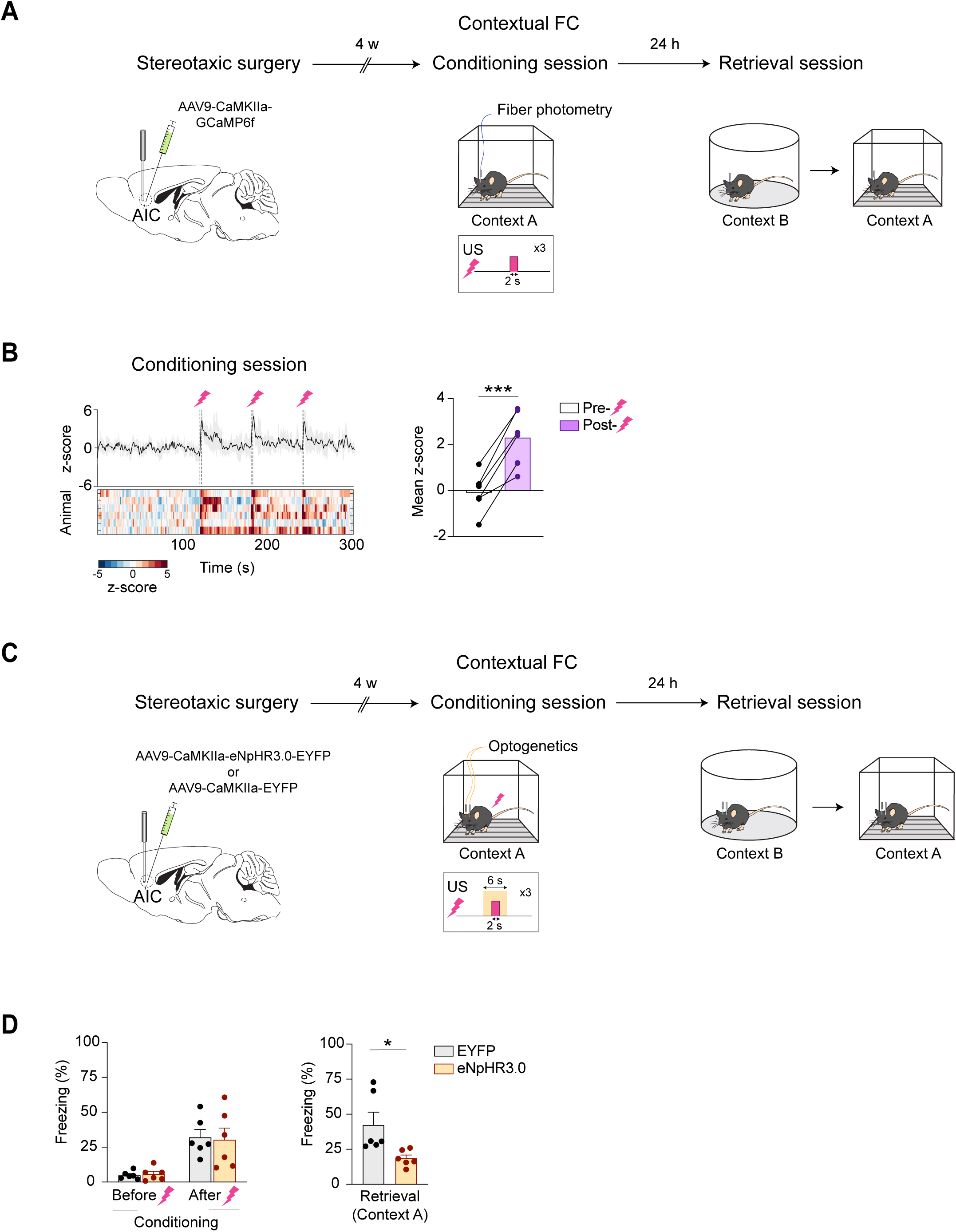
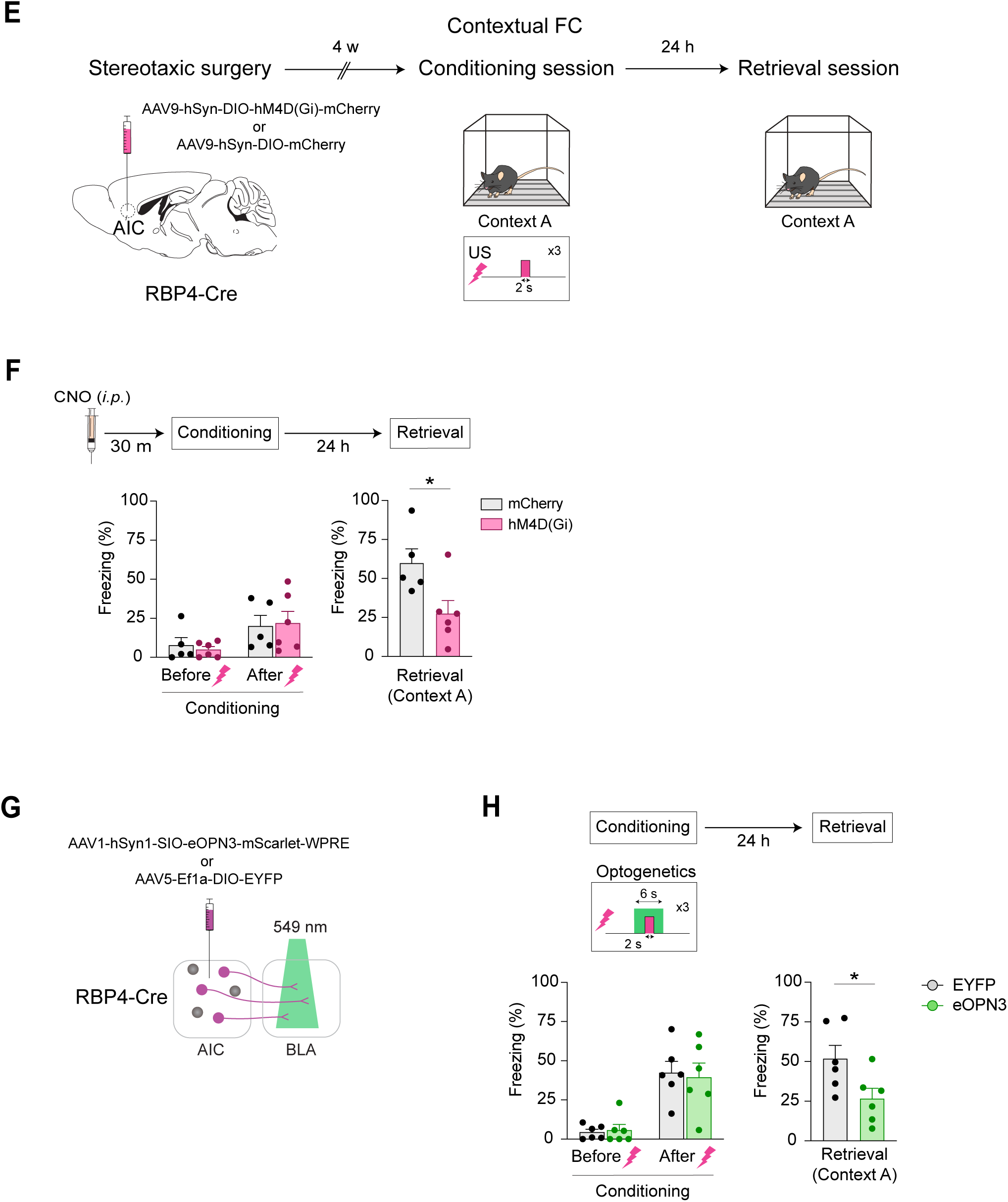
Activity of AIC excitatory neurons, specifically L5-PNs projecting to the BLA, is crucial for the acquisition of contextual fear memory. **(A)** Schematic of the fiber photometry experiments conducted in contextual FC. **(B)** Left: During the conditioning session, population-level activity of AIC excitatory neurons showed robust responses to US delivery across three context-US pairings. Right: Mean activity in the 10-s window following US onset was significantly higher than in the 10-s pre-US baseline window. mean z-score (*N* = 6; Two-tailed paired Student’s *t*-test; *p* = 0.0007). **(C)** Schematic of temporally specific optogenetic inhibition experiments during the conditioning session. **(D)** Optogenetic inhibition of AIC excitatory neurons specifically around each context-US pairing (6-second windows encompassing each 2-second US) during the conditioning session significantly reduced fear responses to the conditioning context (Context A) during the retrieval session. Conditioning (*N*_EYFP=_6, *N*_eNpHR3.0=_6; two-way mixed ANOVA, *p*=0.814). Retrieval (*N*_EYFP=_6, *N*_eNpHR3.0=_6; two-tailed Student’s *t*-test, *p*=0.049). **(E)** Schematic of chemogenetic inhibition experiments targeted specifically to L5-PNs in the AIC. **(F)** In contextual FC, chemogenetic inhibition of AIC L5-PNs during the conditioning session impaired fear memory acquisition. Conditioning (*N*_mCherry=_5, *N*_hM4D(Gi)=_6; two-way mixed ANOVA, *p*=0.636). Retrieval (*N*_mCherry=_5, *N*_hM4D(Gi)=_6; two-tailed Student’s *t*-test, *p*=0.029). **(G)** Schematic of projection-specific optogenetic inhibition experiments during the conditioning or the retrieval session. **(H)** Optogenetic inhibition of the AIC^L5-PNs^→BLA pathway during the time windows surrounding each context-US pairing impaired fear memory acquisition. Conditioning (*N*_EYFP=_6, *N*_eOPN3=_6; two-way mixed ANOVA, *p*=0.711). Retrieval (*N*_EYFP=_6, *N*_eOPN3=_6; two-tailed Student’s *t*-test, *p*=0.039). \**p* < 0.05, \*\*\**p* < 0.001. Data are expressed as mean ± SEM. More detailed statistical information is in Table S1.

Having established that AIC L5-PNs are essential for memory acquisition in trace FC, we next asked whether this neuronal population also contributes to the acquisition of contextual fear memory. *In vivo* fiber photometry recordings from AIC L5-PNs revealed robust US-evoked activity during the conditioning session (**Figures S6A and S6B**), consistent with our recordings from the broader AIC excitatory neuron population (**Figures 6A and 6B**). To determine the causal role of AIC L5-PNs, we chemogenetically inhibited them during the conditioning session in contextual FC. This manipulation significantly reduced fear responses during the retrieval session, indicating that AIC L5-PNs are required for contextual fear memory acquisition (**Figures 6E and 6F**).

Given the critical role of the AIC^L5-PNs^→BLA pathway in fear memory acquisition in trace FC, we next asked whether this pathway is also required for fear memory acquisition in contextual FC. We expressed eOPN3 in AIC L5-PNs and selectively inhibited their terminals in the BLA during the time windows surrounding each context-US pairing (**Figure 6G**). Inhibition of the AIC^L5-PNs^→BLA pathway significantly reduced fear responses during the retrieval session (**Figure 6H**). Together, these findings demonstrate that AIC excitatory neurons, particularly BLA-projecting AIC L5-PNs, are required for fear memory acquisition not only in trace FC but also in contextual FC.

### AIC L5-PNs, projecting to the BLA, constrain fear responses during contextual fear memory retrieval

Our earlier pharmacological AIC inactivation revealed an unexpected role for the AIC in contextual fear memory retrieval, contrasting with its lack of involvement in memory retrieval in trace FC. We therefore sought to further characterize AIC function during contextual fear memory retrieval. First, we examined AIC activity dynamics during this phase. *In vivo* fiber photometry recordings revealed significantly greater population-level activity in AIC excitatory neurons when mice were re-exposed to the conditioning context (Context A) than when they were exposed to a neutral context (Context B) (**Figures 7A and 7B**). Notably, during the retrieval session in trace FC, AIC excitatory neurons also exhibited responses to the CS and TI (**Figures S7A and S7B**), but these retrieval-related activities are not causally required for fear memory retrieval in trace FC, as indicated by our earlier pharmacological data (**Figure 1E**).

**Figure 7.**
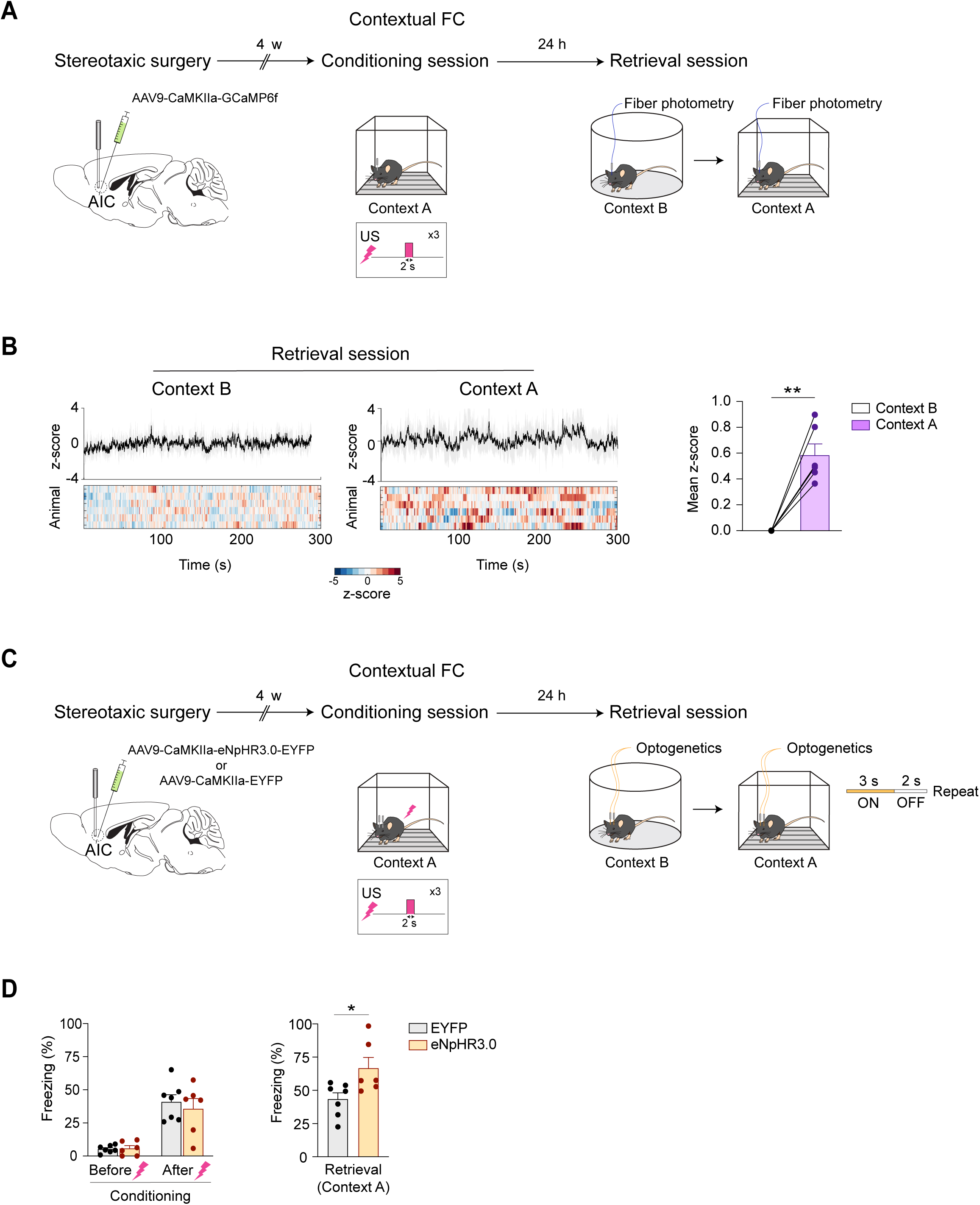
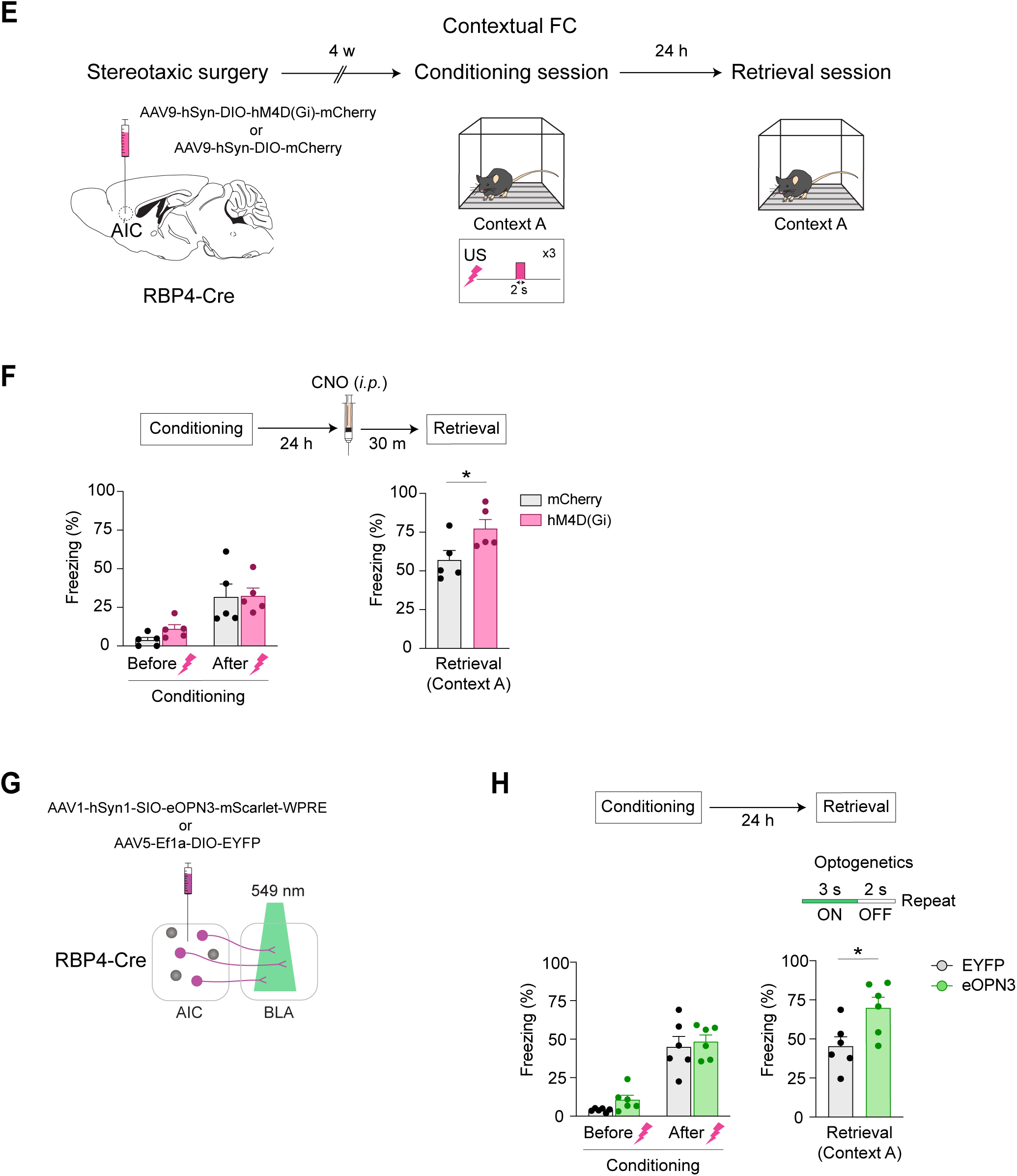
Activity of AIC excitatory neurons, specifically L5-PNs projecting to the BLA, constrain fear responses during contextual fear memory retrieval. **(A)** Schematic of the fiber photometry experiments conducted in contextual FC. **(B)** Left: During the retrieval session, AIC excitatory neurons exhibited elevated, context-driven activity when mice are re-exposed to the conditioning context (Context A), but not when exposed to a neutral context (Context B). Right: Mean activity was significantly higher in Context A than in Context B. mean z-score (*N* = 6; Two-tailed one-sample *t*-test; *p* = 0.0011). **(C)** Schematic of optogenetic inhibition experiments during the retrieval session. **(D)** Optogenetic inhibition of AIC excitatory neurons (3-second ON/2-second OFF cycle) during the retrieval session significantly increased fear responses to the conditioning context (Context A). Conditioning (*N*_EYFP=_7, *N*_eNpHR3.0=_6; two-way mixed ANOVA, *p*=0.476). Retrieval (*N*_EYFP=_7, *N*_eNpHR3.0=_6; two-tailed Student’s *t*-test, *p*=0.026). **(E)** Schematic of chemogenetic inhibition experiments targeted specifically to L5-PNs in the AIC. **(F)** Chemogenetic inhibition of AIC L5-PNs during the retrieval session significantly increased fear responses during memory retrieval. Conditioning (*N*_mCherry=_5, *N*_hM4D(Gi)=_5; two-way mixed ANOVA, *p*=0.485). Retrieval (*N*_mCherry=_5, *N*_hM4D(Gi)=_5; two-tailed Student’s *t*-test, *p*=0.046). **(G)** Schematic of projection-specific optogenetic inhibition experiments during the conditioning or the retrieval session. **(H)** Optogenetic inhibition of the AIC^L5-PNs^→BLA pathway during the retrieval session significantly increased fear responses during memory retrieval. Conditioning (*N*_EYFP=_6, *N*_eOPN3=_6; two-way mixed ANOVA, *p*=0.685). Retrieval (*N*_EYFP=_6, *N*_eOPN3=_6; two-tailed Student’s *t*-test, *p*=0.037). \**p* < 0.05, \*\**p* < 0.01. Data are expressed as mean ± SEM. More detailed statistical information is in Table S1.

We therefore asked whether, in contrast to trace FC, the retrieval-related activity observed during contextual memory retrieval plays a causal role in regulating fear responses. Optogenetic inhibition of AIC excitatory neurons during the retrieval session, using repeated 3-s ON/2-s OFF illumination (**Figure 7C**), significantly increased fear responses in the conditioning context (**Figure 7D**), consistent with the effect of pharmacological AIC inactivation (**Figure 5C**). Importantly, the same manipulation did not increase fear responses in the neutral context (**Figures S8A and S8B**), indicating that the effect was specific to memory retrieval in the threat-associated context rather than reflecting a nonspecific increase in immobility. These findings indicate that AIC excitatory neuron activity normally constrains fear responses during contextual memory retrieval.

We next examined whether AIC L5-PNs also contribute to the regulation of fear responses during contextual fear memory retrieval. *In vivo* fiber photometry recordings from AIC L5-PNs revealed significantly greater activity in the conditioning context than in the neutral context during the retrieval session (**Figures S9A and S9B**), consistent with our recordings from the broader AIC excitatory neuron population (**Figures 7A and 7B**). Chemogenetic inhibition of AIC L5-PNs during the retrieval session significantly increased fear responses in the conditioning context (**Figures 7E and 7F**), demonstrating that AIC L5-PNs causally regulate fear responses during contextual fear memory retrieval. By contrast, although AIC L5-PNs also exhibited CS- and TI-related activity during memory retrieval in trace FC (**Figures S10A and S10B**), our earlier pharmacological data suggest that these responses do not causally regulate fear responses (**Figure 1E**). These findings reveal a paradigm-dependent dissociation in the retrieval-phase function of AIC L5-PNs: context-driven activity regulates fear responses in contextual FC, whereas CS- and TI-related activity does not in trace FC.

Finally, we asked whether the AIC^L5-PNs^→BLA pathway also mediates this regulatory function in the retrieval-phase of contextual FC. Optogenetic inhibition of AIC L5-PN terminals in the BLA during the retrieval session significantly increased fear responses in the conditioning context (**Figures 7G and 7H**). This indicates that the same AIC^L5-PNs^→BLA pathway is also required for regulating fear responses during memory retrieval in a context-dependent manner. Taken together, these findings reveal distinct, phase-specific functions of the AIC: promoting fear memory acquisition, while constraining fear responses during memory retrieval in contextual FC.

### Distinct subpopulations of AIC excitatory neurons are engaged for fear memory acquisition and retrieval

The opposing behavioral effects in contextual FC raise a fundamentally important question: how can the same neuronal population and anatomical pathway mediate such opposing outcomes? We hypothesized that distinct neuronal subpopulations within the AIC^L5-PNs^ are preferentially engaged during specific phases of the process. To test this, we combined ST-Cal-Light, which labels excitatory neurons active during the conditioning session, with c-Fos labeling of neurons active during the retrieval session, focusing specifically on L5 (**Figure 8A**). Notably, AIC L5 neurons active during the conditioning session were largely distinct from those activated during the retrieval session, showing only ∼8% overlap (**Figures 8B and 8C**). This indicates that separable neuronal subpopulations are engaged in memory acquisition versus retrieval. We note that ST-Cal-Light-labeled L5 neurons are specifically excitatory, whereas the c-Fos-labeled population may include a small fraction of inhibitory neurons, although cortical L5 consists predominantly of excitatory pyramidal neurons^56–58^.

**Figure 8.**
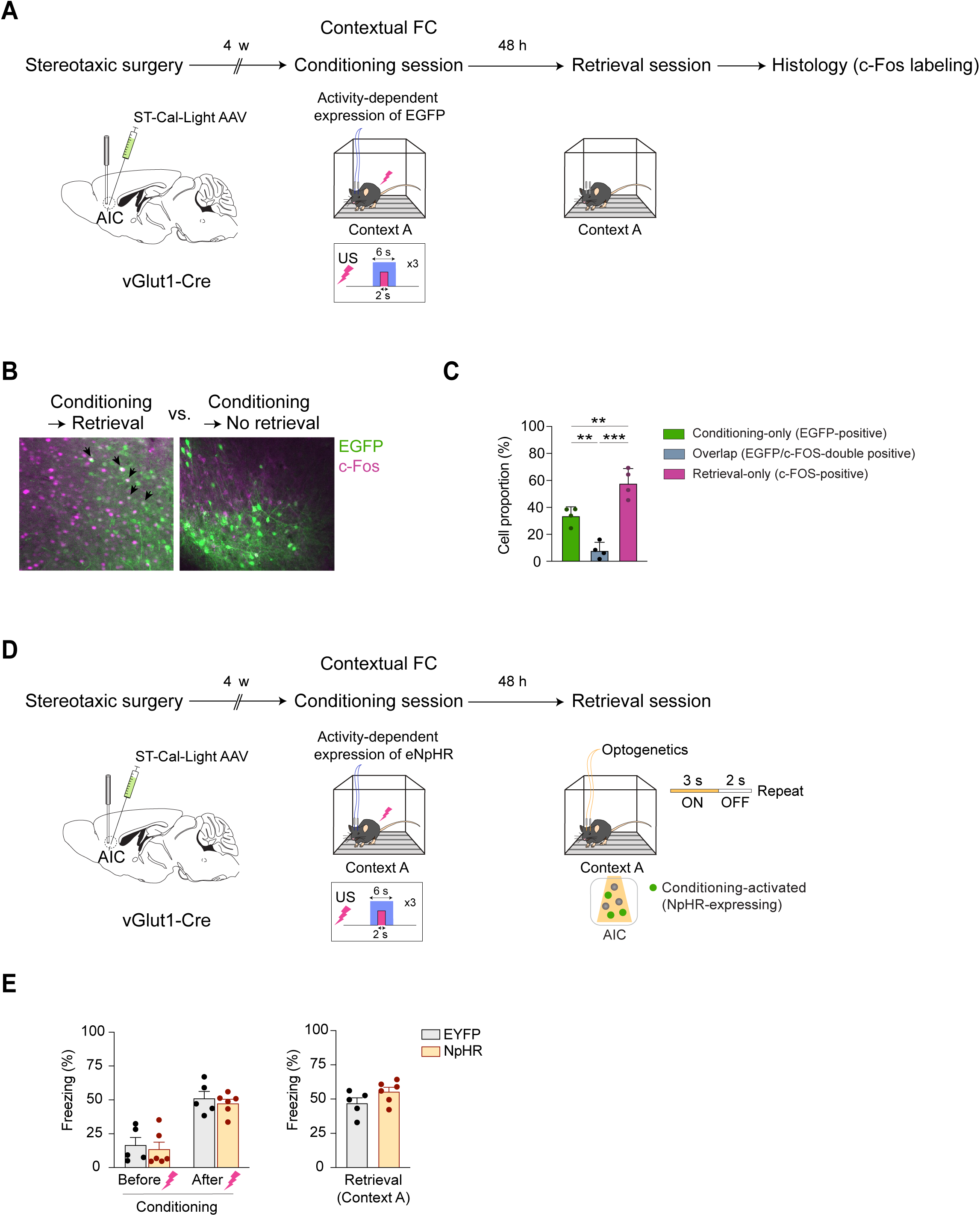
Distinct AIC excitatory neuron ensembles independently mediate fear memory acquisition and its context-dependent retrieval. **(A)** Schematic of the dual ST-Cal-Light and c-Fos labeling approach to capture neuronal populations active during the conditioning session and the retrieval session, respectively. **(B)** Representative images of AIC L5 showing ST-Cal-Light-labeled neurons (green; active during the conditioning session) and c-Fos-positive neurons (magenta; active during the retrieval session). Black arrows indicate neurons double-positive for ST-Cal-Light labeling and c-Fos immunostaining (blue-gray; active during both conditioning and retrieval sessions). Right: A control cohort perfused 48 hours after conditioning without a retrieval session showed minimal c-Fos expression. **(C)** Quantification reveals that AIC L5 neurons activated during the conditioning session and the retrieval session are largely distinct populations, showing only 8% overlap (*N*=4; one-way ANOVA, *p*=3.82×10^-5^; post-hoc Tukey’s HSD). **(D)** Schematic of the optogenetic strategy used to test whether AIC excitatory neurons active during the conditioning session are required for regulating fear responses during the retrieval session. **(E)** Optogenetic inhibition of the conditioning-active neuronal population had no significant effect on fear responses to the conditioning context during the retrieval session. Conditioning (*N*_EYFP=_5, *N*_NpHR=_6; two-way mixed ANOVA, *p*=0.931). Retrieval (*N*_EYFP=_5, *N*_NpHR=_6; two-tailed Student’s *t*-test, *p*=0.139). \*\**p* < 0.01, \*\*\**p* < 0.001. Data are expressed as mean ± SEM. More detailed statistical information is in Table S1.

Next, we investigated the functional specificity of these distinct neuronal subpopulations. We utilized ST-Cal-Light to specifically express the inhibitory opsin NpHR in AIC excitatory neurons that were active during the conditioning session (**Figure 8D**). If the same neuronal subpopulation mediates both acquisition and retrieval, optogenetic inhibition of these conditioning-activated neurons should significantly alter fear responses during the retrieval session. However, inhibiting this neuronal subpopulation during the retrieval session had no significant effect on fear responses in the conditioning context (**Figure 8E**). These results provide functional evidence that the neuronal subpopulation recruited for contextual memory acquisition is largely distinct from that required for retrieval, endorsing the conclusion that largely distinct AIC neuronal subpopulations independently regulate two phases of contextual fear memory.

## DISCUSSION

In this study, we dissected the role of the mouse AIC, particularly its rostral agranular subregion, in fear learning and memory across three distinct FC paradigms (delay, trace, and contextual) and their memory phases (acquisition, consolidation, and retrieval), respectively. By integrating pharmacological, chemogenetic, optogenetic, fiber photometry, and activity-dependent labeling approaches, we found that AIC excitatory neurons, particularly L5-PNs that project to the BLA, play a crucial role in fear memory acquisition in trace and contextual FC but not in delay FC, suggesting their selective engagement under cognitively demanding conditions. In addition to this primary novel finding on the higher-order functions of the AIC in fear learning and memory, we also observed that, only in contextual FC but not in trace FC, a separate subset of AIC L5-PNs distinct from those required for memory acquisition also plays a role in memory retrieval. Because our manipulations and recordings specifically targeted the rostral AIC, where the insular cortex is exclusively agranular, these findings define functions of the rostral agranular AIC rather than those of the more caudal AIC, which spans multiple cytoarchitectonic subdivisions and has been the focus of previous studies. Altogether, we provide new mechanistic and conceptual insights into the paradigm- and memory phase-specific higher-order functions of the rostral agranular AIC in fear learning and memory.

### Common involvement of the AIC for fear memory acquisition in both trace and contextual FC

Our results reveal that AIC excitatory neurons, particularly L5-PNs, exhibit an emergent and transient activity precisely time-locked to the onset of the TI during trace FC, which plays a pivotal role in fear memory acquisition. This temporally precise, transient activity may serve as a signal that triggers downstream networks to maintain CS information over the TI. This type of mechanism is analogous to findings in working memory research, where brief neuronal activations are thought to initiate or modulate more sustained activity in distributed circuits that maintain information over delays^59–63^. The AIC is known to evaluate emotional significance by integrating interoceptive signals with exteroceptive input^33,34^. Thus, the TI-related activity we observed might reflect the integration of the CS-associated emotional salience with the temporal discontiguity introduced by the TI. Such integration may help the brain generate a predictive representation of the forthcoming aversive event despite the loss of direct temporal contiguity between CS and US. Consistent with this interpretation, selective inhibition of AIC L5-PNs during the TI impaired fear responses to the CS during the retrieval session, without affecting pre-CS baseline fear responses, indicating a specific disruption of the CS-US association rather than a nonspecific alteration in fear or motor behavior.

This framework may also extend to contextual FC. Importantly, the TI in trace FC is not an information-free interval: although the discrete CS has ended and the US has not yet occurred, the surrounding context remains continuously the same. Previous studies have suggested that contextual information may become significant in fear memory processing when a temporal gap exists between the CS and US^18,64^.

Therefore, TI-related AIC activity may help link the fading representation of the CS with the ongoing context. Altogether, context may therefore provide a common associative substrate across trace and contextual FC. From this perspective, AIC L5-PNs may perform a common function across both paradigms by supporting aversive learning when the US cannot be associated through direct temporal pairing with a discrete, contiguous cue.

### Dual roles of the AIC during memory acquisition and retrieval in contextual FC

Beyond the shared mechanism of the AIC for memory acquisition in trace and contextual FC, we also observed a paradoxical phenomenon: silencing AIC excitatory neurons during contextual retrieval significantly increased fear responses. These data suggest that the AIC may also act to maintain an appropriate level of defensive behavior in a context-dependent manner.

Our data further showed that both functions are mediated by the AIC^L5-PNs^→BLA pathway. This raises an important question: how can the same pathway promote fear memory acquisition yet constrain fear responses during retrieval? Our activity-dependent labeling provided a possible explanation. AIC L5-PNs recruited during acquisition and retrieval formed largely non-overlapping ensembles, and selective inhibition of the acquisition-recruited ensemble had no effect on retrieval. These findings suggest that distinct neuronal ensembles within the same pathway mediate its opposing functions in a memory phase-specific manner. These ensembles may differentially engage functionally distinct BLA microcircuits^65–67^. Specifically, the acquisition-recruited ensemble may preferentially engage fear-promoting BLA neurons underlying fear memory acquisition, whereas the retrieval-recruited ensemble may recruit fear-suppressing BLA neurons or local inhibitory circuits, such as intercalated cells, thereby constraining fear responses during retrieval.

### Future perspectives

Although the following topics are out of scope of the present study, there are at least three potential avenues for future investigations. While our study focused on defining the functional contribution of the top-down AIC^L5-PNs^→BLA pathway, our anatomical data suggest that this pathway likely operates within a broader reciprocal circuit. Retrograde viral tracing revealed robust labeling of BLA neurons projecting to the AIC (**Figures S11A and S11B**), and BLA axons formed dense terminal fields in AIC L5 (**Figure S11C**). Consistent with these findings, cell type-specific monosynaptic rabies tracing confirmed that AIC L5-PNs receive direct synaptic input from BLA neurons (**Figure S11D**). Accordingly, exploring how reciprocal AIC-BLA interactions contribute to higher-order threat learning represents an outstanding question for future investigation.

Second, anatomical and functional differences between the AIC and PIC are a frontline topic^33,34^. Indeed, our study demonstrated that the PIC was not involved in memory acquisition in trace FC. A similar study design to the present investigation that comprehensively examines the role of the insular cortex in higher-order functions of fear learning and memory may be useful to further characterize the differences between the AIC and PIC.

Lastly, it is important to investigate these higher-order functions of the AIC in fear learning and memory from clinical and pathological perspectives. To this end, a transdiagnostic, dimensional framework, such as the National Institute of Mental Health’s Research Domain Criteria (RDoC), may provide a useful approach for mapping AIC circuit dysfunctions onto continuous pathophysiological dimensions that cut across traditional categorical boundaries^8,68,69^. Rather than being restricted to PTSD and anxiety-related diagnoses, alterations in AIC-dependent temporal and contextual processing may represent a shared neurobiological vulnerability underlying abnormalities in threat and safety processing across multiple neuropsychiatric conditions. Indeed, AIC dysfunction has been implicated across a broad range of neuropsychiatric conditions, including schizophrenia and mood disorders, suggesting that it may contribute to symptomatic dimensions shared across diagnostic categories^33,70,71^. Stratifying clinical populations based on such circuit-defined dimensions may therefore facilitate the development of mechanism-based therapeutics and more personalized clinical interventions.

## STAR METHODS

Detailed methods are provided in the online version of this paper and include the following:

- KEY RESOURCES TABLE
- RESOURCE AVAILABILITY

- Lead Contact
- Materials Availability
- Data and Code Availability
- EXPERIMENTAL MODEL AND SUBJECT DETAILS

- Animals
- METHOD DETAILS

- Viral constructs and stereotaxic surgeries
- Pharmacological experiments
- Fear conditioning
- Fiber photometry
- Optogenetics
- Chemogenetics
- Activity-dependent cell labeling
- Histology
- QUANTIFICATION AND STATISTICAL ANALYSIS

## SUPPLEMENTAL INFORMATION

Supplemental Information can be found online

## ACKNOWLEDGMENTS

We thank Ms. Yukiko Lema for organizing the manuscript and figures. We are grateful to Drs. Patricia Janak, Seung-Chan Lee, and John Issa for their insightful feedback on the manuscript. We also thank Dr. Solange Brown for providing the RBP4-Cre mice and acknowledge Victoria Stepanyants and Dr. Brian Lee for conducting a pilot electrophysiological study. Additionally, we thank Drs. Kenichiro Nagahama and Jung Ho Hyun for valuable discussions regarding the ST-Cal-Light experiments. H.N. gratefully acknowledges fellowship fundings from the American Academy of Neurology and American Heart Association. This research was supported by grants from the National Institutes of Health (P50MH136297 to A.S.; DP1MH119428 to H.-B.K.).

## AUTHOR CONTRIBUTIONS

H.N. formulated the conceptual framework, designed and conducted experiments, and wrote the manuscript. B.M., J.C., S.L., L.G., I.I., I.M., A.Saito., K.I., conducted experiments and/or contributed to relevant discussions. H-B.K. assisted in formulating the conceptual framework and designing the neurobiological experiments. A.S. formulated the conceptual framework, supervised the overall project, and wrote the manuscript.

## STAR METHODS

### KEY RESOURCES TABLE

| REAGENT or RESOURCE | SOURCE | IDENTIFIER |
| --- | --- | --- |
| **Antibodies** |  |  |
| Anti-GFP Antibody | Proteintech | 3h9-20 |
| Anti-RFP Antibody | Proteintech | 5f8-20 |
| Anti-c-Fos Antibody | Cell Signaling Technology | 2250T |
| **Bacterial and Virus Strains** |  |  |
| AAV9-CaMKII-GCaMP6f-WPRE-SV40 | Addgene | 100834-AAV9 |
| AAV9-Syn-Flex-GCaMP6f-WPRE-SV40 | Addgene | 100833-AAV9 |
| AAV9-CaMKIIa- eNpHR3.0-EYFP | Addgene | 26971-AAV9 |
| AAV5-CaMKIIa-EYFP | Addgene | 50469-AAV5 |
| AAV1-hSyn1-SIO-eOPN3-mScarlet-WPRE | Addgene | 125713-AAV1 |
| AAV5-Ef1a-DIO-EYFP | Addgene | 27056-AAV5 |
| AAV1-CaMKIIa- eNpHR3.0-EYFP | Addgene | 26971-AAV1 |
| AAV5-Ef1a-DIO-eNpHR3.0-EYFP | Addgene | 26966-AAV5 |
| AAV9-CaMKIIa-hM4D(Gi)-mCherry | Addgene | 50477-AAV9 |
| AAV9-CaMKIIa-mCherry | Addgene | 114469-AAV9 |
| AAV9-hSyn-DIO-hM4D(Gi)-mCherry | Addgene | 44362-AAV9 |
| AAV9-hSyn-DIO-mCherry | Addgene | 50459-AAV9 |
| AAV1-CAG-Flex-EGFP-WPRE-bGH | Addgene | 51502-AAV1 |
| rAAV2-retro CAG-GFP | Addgene | 37825-AAVrg |
| AAV8-Flex-RG | UNC Vector core | N/A |
| AAV8-Flex-TVAmCherry | UNC Vector core | N/A |
| RV-SADΔG-EGFP (EnvA) | Salk Viral Vector Core | N/A |
| AAV-CMV-IE94-Flex-TM-KA2-CaM-NES-TeV-N-AsLOV2-TEVseq-bGH-PA | Addgene | 171619 |
| AAV-hSYN-M13-TEV-C-P2A-tdTomato | Addgene | 92391 |
| AAV-TRE-EGFP | Addgene | 89875 |
| AAV-TRE-NpHR-EYFP | Addgene | 171621 |
| **Chemicals, Peptides, and Recombinant Proteins** |  |  |
| Muscimol | TOCRIS | 028945 |
| Baclofen | TOCRIS | 0417 |
| Clozapine N-oxide | TOCRIS | 6329 |
| **Experimental Models: Organisms/Strains** |  |  |
| C57BL/6J | The Jackson Laboratory | 000664 |
| RBP4-Cre mice | MMRRC | 031125-UCD |
| vGlut1-Cre mice | The Jackson Laboratory | 023527 |
| **Software and Algorithms** |  |  |
| Matlab | Mathworks | R2019a |
| SPSS statistics | IBM | Version 26 |

### RESOURCE AVAILABILITY

#### Lead Contact

Further information and requests for resources and reagents should be directed to and will be fulfilled by the Lead Contact, Akira Sawa.

#### Materials Availability

Materials related to this study are available from the Lead Contact on reasonable request.

#### Data and Code Availability

All major data generated or analyzed are included in the article. Any additional data related to this study are available from the Lead Contact upon reasonable request. No unique code was generated for this study; instead, it utilized previously established algorithms and codes, as described in the **METHOD DETAILS**.

### EXPERIMENTAL MODEL AND SUBJECT DETAILS

#### Animals

Adult male mice aged 9-18 weeks were used in this study. Wild-type C57BL/6J mice (Jackson Laboratory) were employed for pharmacological inactivation, chemogenetic and optogenetic inhibition, and fiber photometry. RBP4-Cre mice^54^ on the C57BL/6J background, which have been shown to express Cre recombinase in cortical layer 5 pyramidal neurons^72^, were utilized for cell type-specific optogenetic and chemogenetic inhibition, pathway-specific optogenetic inhibition, fiber photometry, and viral circuit tracing. vGlut1-Cre mice on a C57BL/6J background were used for activity-dependent cell labeling with ST-Cal-Light. All experimental procedures were conducted in accordance with NIH guidelines and approved by the Johns Hopkins Animal Care and Use Committee.

### METHOD DETAILS

#### Viral constructs and stereotaxic surgeries

For fiber photometry recordings of AIC excitatory neurons, wild-type mice received a unilateral injection of AAV9-CaMKII-GCaMP6f-WPRE-SV40 (0.5 μL; catalog #100834-AAV9, Addgene) into the AIC at coordinates relative to Bregma: AP +2.34 mm, ML −2.3 mm, DV −2.7 mm. A fiber optic cannula (MFC_400-430-0.48_5mm, Doric Lenses) was then implanted at the same site and secured to the skull with acrylic glue. For optogenetic inhibition of AIC excitatory neurons, mice were bilaterally injected with either AAV9-CaMKIIa-eNpHR3.0-EYFP (0.5 μL; catalog #26971-AAV9, Addgene) or, as a control, AAV5-CaMKIIa-EYFP (0.5 μL; catalog #50469-AAV5, Addgene) into the AIC at coordinates relative to Bregma: AP +2.34 mm, ML ±2.3 mm, DV −2.7 mm. Fiber optic cannulas were implanted 0.5 mm above the injection sites and secured with acrylic glue. For optogenetic inhibition of AIC L5-PNs projecting to either the BLA or dmPFC, RBP4-Cre mice were bilaterally injected with either AAV1-hSyn1-SIO-eOPN3-mScarlet-WPRE (0.5 μL; catalog #125713-AAV1, Addgene) or, as a control, AAV5-Ef1a-DIO-EYFP (0.5 μL; catalog #27056-AAV5, Addgene) into the AIC using the same coordinates as above. To inhibit axonal terminals in either the BLA or dmPFC, fiber optic cannulas were bilaterally implanted at the following coordinates and secured with acrylic glue: BLA (AP −1.22 mm, ML ±3.27 mm, DV −4.2 mm) and dmPFC (AP +1.94 mm, ML ±0.5 mm, DV −1.0 mm). For chemogenetic inhibition of AIC excitatory neurons, wild-type mice were bilaterally injected into the AIC (using the coordinates described above) with either AAV9-CaMKIIa-hM4D(Gi)-mCherry (0.5 μL; catalog #50477-AAV9, Addgene) or, as a control, AAV9-CaMKIIa-mCherry (0.5 μL; catalog #114469-AAV9, Addgene). For chemogenetic inhibition of AIC L5-PNs, RBP4-Cre mice were bilaterally injected into the AIC with either AAV9-hSyn-DIO-hM4D(Gi)-mCherry (0.5 μL; catalog #44362-AAV9, Addgene) or, as a control, AAV9-hSyn-DIO-mCherry (0.5 μL; catalog #50459-AAV9, Addgene) using the same coordinates. For anterograde tracing of AIC L5-PNs, RBP4-Cre mice were unilaterally injected with AAV1-CAG-Flex-EGFP-WPRE-bGH (0.5 μL; catalog #51502-AAV1, Addgene). For retrograde tracing of afferent inputs to the AIC, a unilateral injection of rAAV2-retro CAG-GFP (0.5 μL; catalog #37825-AAVrg, Addgene) was administered into the AIC at coordinates: AP +2.34 mm, ML −2.3 mm, DV −2.7 mm. For monosynaptic retrograde rabies tracing of AIC L5-PNs, RBP4-Cre mice were first unilaterally injected with 0.5 μL of a 6:1 (RG:TVA) mixture of helper viruses—AAV8-Flex-RG (UNC Vector Core) and AAV8-Flex-TVAmCherry (UNC Vector Core)—into the AIC at coordinates: AP +2.34 mm, ML −2.3 mm, DV −2.7 mm. Three weeks later, 0.5 μL of RV-SADΔG-EGFP (EnvA; Salk Viral Vector Core) was injected at the same coordinate, and mice were perfused for histological analysis 7-10 days after the rabies virus injection. For all viral injections, mice were deeply anesthetized via an intraperitoneal injection of Avertin (0.4 mg/g) and positioned in a stereotaxic apparatus (David Kopf Instruments). The head was leveled using the Bregma and Lambda reference points, and a craniotomy was performed. Viruses were injected using a Hamilton syringe (catalog #80383); the syringe was left in place for 5 minutes after each injection before being slowly withdrawn. The incision was sutured, and mice were allowed to recover from anesthesia on a heating pad before being returned to their home cages. Animals were given at least 3 weeks of recovery following surgery, except for tracing experiments.

#### Fear conditioning

Delay and trace fear conditioning (FC) paradigms were employed in a comparative manner. Each paradigm consisted of three experimental phases (habituation, fear acquisition, and retrieval) conducted on three consecutive days in large, soundproof isolation chambers (Med Associates Inc.). Mice underwent a 10-minute habituation phase in a conditioning context one day before the conditioning session. On Day 1, mice in the conditioning context were trained to associate a pure tone (conditioned stimulus, CS; 3 kHz, 70 dB, 20 s) with a foot shock (unconditioned stimulus, US; 2 s, 0.5 mA) through four repeated CS-US pairings, each separated by a variable interval (160-200 s). Conditioning performance (learning curve) was assessed by recording and averaging freezing responses during the four CS presentations. Unless otherwise noted, freezing behavior was automatically scored using VideoFreeze™ Video Fear Conditioning Software (Med Associates Inc.). A baseline (BL) response was defined as the mean percentage of freezing during the initial 2-minute period before the first CS presentation. The two paradigms differed only in the interval between the CS and US: in delay FC, the 2-s US co-terminated with the 20-s CS, whereas in trace FC an 18-s interval (the trace interval) was inserted between the termination of the 20-s CS and the onset of the 2-s US. On Day 2 (24 hours after the conditioning session), cued fear retrieval was evaluated by recording and averaging freezing responses during four CS presentations in a novel context distinct from the conditioning context, with each CS presented at a variable interval (160-200 s). A baseline (BL) response was measured as the mean percentage of freezing during the initial 2-minute period preceding the first CS presentation. Contextual FC also comprised three experimental phases (habituation, fear acquisition, and retrieval) conducted over three consecutive days in the same isolation chambers. Mice were habituated for 10 minutes in a conditioning context one day before the conditioning session. On Day 1, in the conditioning context, mice received three foot shocks (US; 2 s, 0.5 mA) delivered at 2, 3, and 4 minutes during a 5-minute session, without any discrete cue as used in the cued FC protocol. Freezing responses were measured as the mean percentage of time spent freezing during the first 2 minutes (pre-shock) and the last 3 minutes (post-shock) of the session. On Day 2, contextual fear retrieval was assessed over 5 minutes by recording freezing responses in either the conditioning context (Context A) or a novel context (Context B). In each context, freezing responses were calculated as the mean percentage of time spent freezing during the 5-minute period. For mice tethered to fiber-optic patch cords during optogenetic experiments, freezing behavior was instead analyzed using ezTrack’s Freeze Analysis Module^73^, which allowed us to minimize artifacts caused by cable movement that could otherwise be misclassified as animal motion.

#### Pharmacological experiments

For drug infusions, mice were bilaterally implanted with 26-gauge guide cannulas (8IC315GSPCXC, PlasticsOne) targeting the AIC at the following coordinates relative to Bregma: AP +2.34 mm, ML ±2.3 mm, DV −2.7 mm, and secured to the skull with acrylic glue. Intracranial microinjections were performed using a MICRO-4 microsyringe pump controller (World Precision Instruments) connected to a Hamilton microsyringe (PlasticsOne, HSYR-1) via a tube (PlasticsOne, C230C/SPC) at a flow rate of 0.15 μL/min. For pharmacological inactivation of the AIC, a solution containing Muscimol (0.5 μg) and Baclofen (0.5 μg) in 0.3 μL per side (or saline as a control) was bilaterally infused into the AIC 30 minutes before FC or prior to retrieval, depending on the experimental goal. To approximate the cannula tip location and the drug diffusion area, a fluorescent dye was infused after all experimental sessions were completed, thereby avoiding any potential chemical or volumetric interactions with the Muscimol/Baclofen solution.

#### Fiber photometry

Fiber photometry was employed to monitor Ca²⁺-sensitive fluorescence in genetically defined cell populations. Two LEDs, 465 nm for GCaMP6f excitation and 405 nm for isosbestic excitation (CLED_465 and CLED_405, Doric Lenses), were synchronized using a TTL pulse generator (OTPG4, Doric Lenses) to alternate excitation at 10 Hz. Excitation light was coupled into a low-autofluorescence 400 μm patch cord (0.48 NA, 1 m cable length, Doric Lenses), with one end connected via a ferrule (SLEEVE_ZR_1.25-BK, Doric Lenses) to a fiber optic cannula (MFC_400-430-0.48_5mm, Doric Lenses) implanted into the AIC. The LED power for both wavelengths was measured with a power meter (PM100D, Thorlabs) and adjusted to 50-80 μW at the tip of the patch cord. Emission light from GCaMP6f was collected through the same fiber and detected using an sCMOS camera (Zyla 4.2, Andor), which was synchronized with the LEDs via the same TTL pulse generator to capture signals at 20 Hz and controlled by Micro-Manager software. To calculate ΔF/F, the 405 nm signal was fit to the 465 nm signal. This produced a fitted 405 nm signal used to normalize the 465 nm signal as follows: ΔF/F = (465 nm signal − fitted 405 nm signal) / fitted 405 nm signal. Event-related changes in calcium signals were further normalized using z-scoring. For the trace FC paradigm, activity during the CS-TI-US period was quantified by calculating a z-score based on the mean and standard deviation (SD) of the 30 s baseline preceding each CS onset. For the contextual FC paradigm, US-evoked responses were z-scored using the mean and SD of the 2-min baseline prior to the first US delivery. To assess activity during re-exposure to the conditioning context (context A), z-scores were calculated using the mean and SD of the activity recorded in the neutral context (context B).

#### Optogenetics

Optogenetics was employed to manipulate neural activity in genetically defined cell types within the AIC and its connected pathways. For optogenetic inhibition, mice were bilaterally implanted with fiber optic cannulas (MFC_200-245-0.53_5mm_ZF1.25_FLT, Doric Lenses). During experiments, these implanted cannulas were connected via ferrules (SLEEVE_ZR_1.25-BK, Doric Lenses) to the bifurcated ends of a branching fiber-optic patch cord that was pre-connected to a fiber-optic rotary joint (FRJ_1x1_FC-FC, Doric Lenses) to allow free movement. Another patch cord was connected from the rotary joint to either a 612 nm laser (Ce:YAG & Blue Fiber Light Source/CF-Custom Filter FF01-612/69, Doric Lenses) or a 549 nm laser (Ce:YAG Fiber Light Source/CF-Custom Filter FF01-549/15, Doric Lenses) for continuous stimulation of eNpHR3.0 or eOPN3, respectively. The laser power for both wavelengths was measured with a power meter (PM100D, Thorlabs) and set to 10 mW at the tip. Laser power was verified before and after every FC session to ensure proper equipment function. Laser output was synchronized by a TTL pulse generator (OTPG4, Doric Lenses) and controlled by Doric Neuroscience Studio software (Doric Lenses).

#### Chemogenetics

Clozapine N-oxide (CNO; Tocris, catalog #6329) was dissolved in saline (3 mg/kg) and administered via intraperitoneal (i.p.) injection.

#### Activity-dependent cell labeling

A recently developed tool, soma-targeted Cal-Light (ST-Cal-Light)^53^, was used to label AIC excitatory neurons active within defined time windows. AAVs encoding ST-KA2, M13-TEV-C, and a TRE-driven effector were mixed at a 2:2:5 ratio (AAV-pCMV-FLEX-TM-KA2-CaM-NES-TEV-N-AsLOV2-TEVseq-tTA : AAV-hSYN-M13-TEV-C-P2A-tdTomato : AAV-TRE-EGFP **or** AAV-TRE-NpHR-EYFP; total volume 0.5 μl) and injected into the AIC of vGlut1-Cre mice (relative to bregma: AP +2.34 mm, ML −2.30 mm, DV −2.70 mm). For optical labeling, blue light (473 nm) was delivered to the AIC during designated time windows. Power was measured at the fiber tip and adjusted to 5-10 mW. Brains were collected for histological analysis 48 h after the labeling experiments.

#### Histology

Mice were euthanized and transcardially perfused with PBS, followed immediately by 4% paraformaldehyde (PFA). Brains were post-fixed overnight in 4% PFA at 4°C and then coronally sectioned at 50 µm using a Leica cryostat. Brain sections were incubated for 16 hours at 4°C with primary antibodies diluted in a blocking solution of 10% normal goat serum (NGS) in PBST (PBS with 1% BSA and 0.5% Triton X-100). Primary antibodies used were: anti-GFP (1:500; clone 3h9-20, Proteintech), anti-RFP (1:500; clone 5f8-20, Proteintech), and anti-c-fos (1:1000; 2250T, Cell Signaling Technology). The sections were then washed and incubated for 2 hours at room temperature with the corresponding secondary antibodies in 1% NGS in PBST: Alexa Fluor 488 goat anti-rat (1:400; Invitrogen, A11006), Alexa Fluor 568 goat anti-rat (1:400; Invitrogen, A11077), or Alexa Fluor 647 goat anti-rabbit (1:500; Invitrogen). After washing, sections were stained with DAPI (Invitrogen). All sections were mounted and imaged using either a Zeiss AX10 Observer D1 or a Zeiss LSM800 confocal microscope at the Johns Hopkins University Microscope Facility.

### QUANTIFICATION AND STATISTICAL ANALYSIS

Detailed statistical information for each figure is summarized in **Table S1**, complementing the concise descriptions provided in the corresponding figure legends. Briefly, no formal statistical methods were used to predetermine sample sizes; however, our sample sizes are consistent with those commonly employed in the field. For parametric tests such as ANOVAs and t-tests, data were assumed to follow a normal distribution given that these tests are robust to deviations from normality. For analyses involving one within-subjects factor and one between-subjects factor, a two-way mixed ANOVA was performed with the Greenhouse-Geisser correction applied as necessary to account for violations of the sphericity assumption. For analyses involving one repeated measure, a one-way repeated measures ANOVA was used, while for analyses involving one non-repeated factor, a one-way ANOVA followed by post hoc Tukey’s HSD test was applied. For within-subject comparisons, a two-tailed paired Student’s t-test was utilized. For a comparison of independent two groups, two-tailed Student’s or Welch’s t-test was used, depending on the homogeneity of variance as assessed by Levene’s test.

