## Supplementary information for "The anterior insular cortex mediates higher-order threat learning and memory"

### Fig. S1

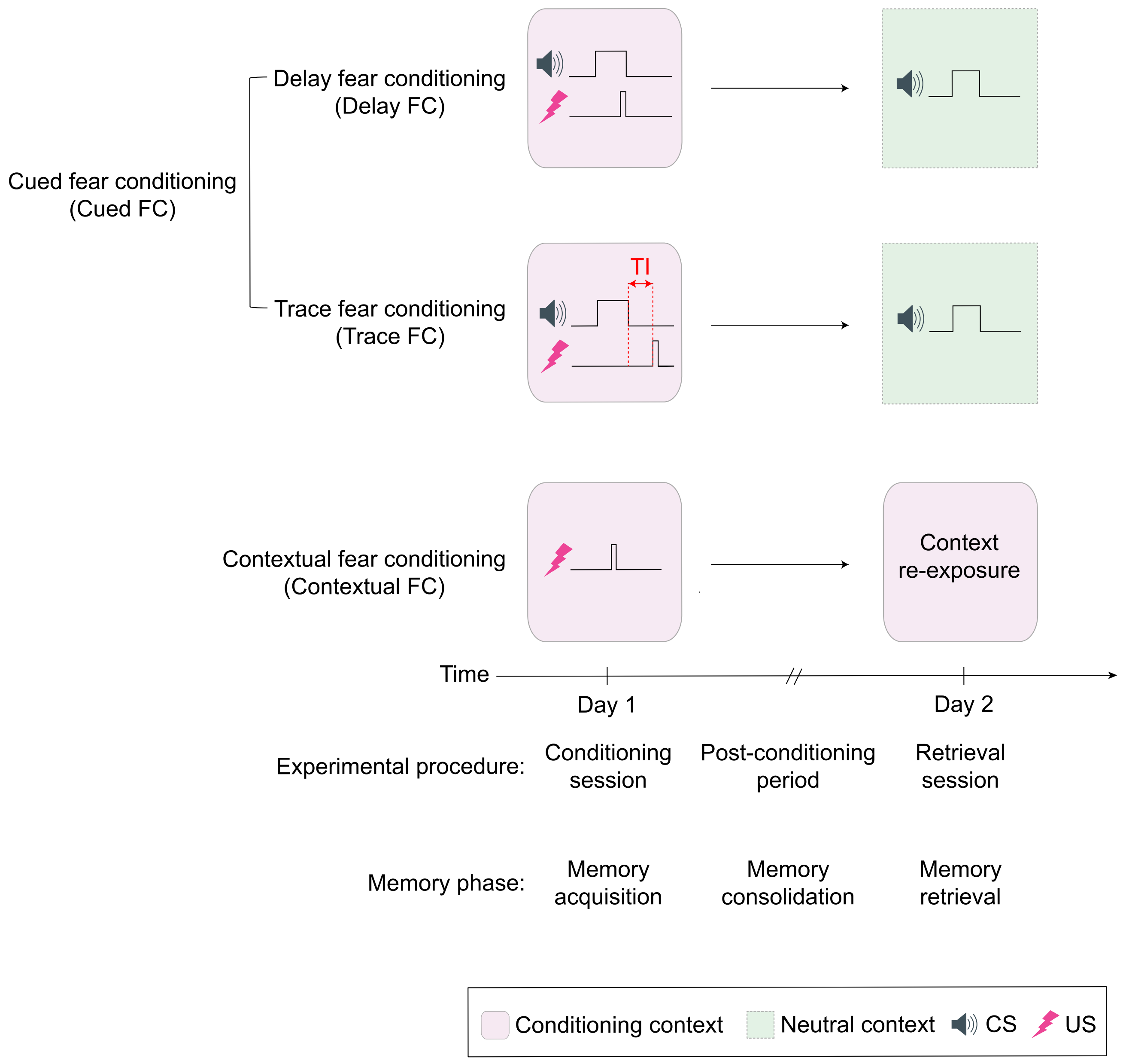

Cued fear conditioning
(Cued FC)
Delay fear conditioning
(Delay FC)
Trace fear conditioning
(Trace FC)
TI
Contextual fear conditioning
(Contextual FC)
Context
re-exposure
Time
Day 1
Day 2
Experimental procedure:
Conditioning
session
Post-conditioning
period
Retrieval
session
Memory phase:
Memory
acquisition
Memory
consolidation
Memory
retrieval
Conditioning context
Neutral context
CS
US

**Figure S1. Schematic overview of fear conditioning paradigms and corresponding memory phases.** Fear conditioning (FC) paradigms are broadly categorized into cued and contextual FC. Cued FC is further divided into delay and trace FC. In delay FC, the CS and US are temporally contiguous, whereas in trace FC, the CS and US are separated by a trace interval (TI). In contextual FC, the US is associated with the conditioning context rather than a discrete CS. Across paradigms, the conditioning session corresponds to memory acquisition, the post-conditioning period to memory consolidation, and the retrieval session to memory retrieval. During retrieval, cued fear memory is assessed by CS presentation in a neutral context, whereas contextual fear memory is assessed by re-exposure to the conditioning context. CS, conditioned stimulus; US, unconditioned stimulus; TI, trace interval.

### Fig. S2

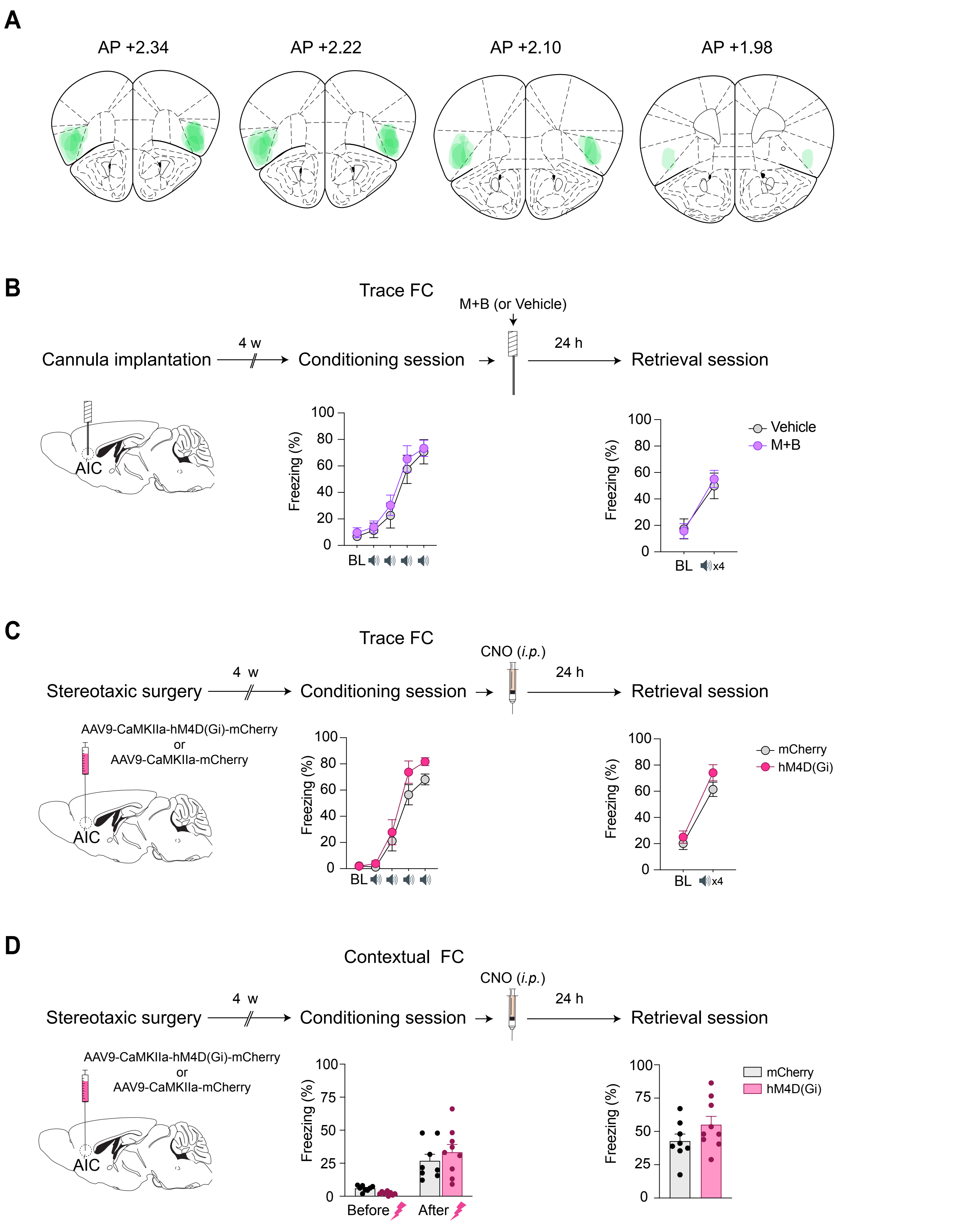

A
AP +2.34
AP +2.22
AP +2.10
AP +1.98
B
Trace FC
M+B (or Vehicle)
Cannula implantation
4 w
Conditioning session
24 h
Retrieval session
AIC
Freezing (%)
BL
x4
Vehicle
M+B
C
Trace FC
CNO (i.p.)
Stereotaxic surgery
4 w
Conditioning session
24 h
Retrieval session
AAV9-CaMKIIa-hM4D(Gi)-mCherry
or
AAV9-CaMKIIa-mCherry
AIC
Freezing (%)
BL
x4
mCherry
hM4D(Gi)
D
Contextual FC
CNO (i.p.)
Stereotaxic surgery
4 w
Conditioning session
24 h
Retrieval session
AAV9-CaMKIIa-hM4D(Gi)-mCherry
or
AAV9-CaMKIIa-mCherry
AIC
Freezing (%)
Before
After
mCherry
hM4D(Gi)

**Figure S2. AIC activity is not required for fear memory consolidation in either trace or contextual FC.**

**(A)** Coronal schematics illustrating the estimated extent of drug diffusion. It appears that the drug was primarily infused into the rostral part of the AIC, the subregion characterized by agranular cytoarchitecture.

**(B)** In trace FC, pharmacological inactivation of the AIC during the post-conditioning period did not disrupt fear memory acquisition. Conditioning ($N_{Vehicle}$=9, $N_{M+B}$=9; two-way mixed ANOVA, $p$=0.922). Retrieval ($N_{Vehicle}$=9, $N_{M+B}$=9; two-way mixed ANOVA, $p$=0.587).

**(C)** In trace FC, chemogenetic inhibition of AIC excitatory neurons during the post-conditioning period did not impact fear memory acquisition. Conditioning ($N_{mCherry}$=9, $N_{hM4D(Gi)}$=9; two-way mixed ANOVA, $p$=0.326). Retrieval ($N_{mCherry}$=9, $N_{hM4D(Gi)}$=9; two-way mixed ANOVA, $p$=0.441).

**(D)** In contextual FC, chemogenetic inhibition of AIC excitatory neurons during the post-conditioning period did not affect fear memory acquisition. Conditioning ($N_{mCherry}$=8, $N_{hM4D(Gi)}$=9; two-way mixed ANOVA, $p$=0.212). Retrieval ($N_{mCherry}$=8, $N_{hM4D(Gi)}$=9; two-tailed Student's $t$-test, $p$=0.156).

More detailed statistical information is in Table S1.

### Fig. S3

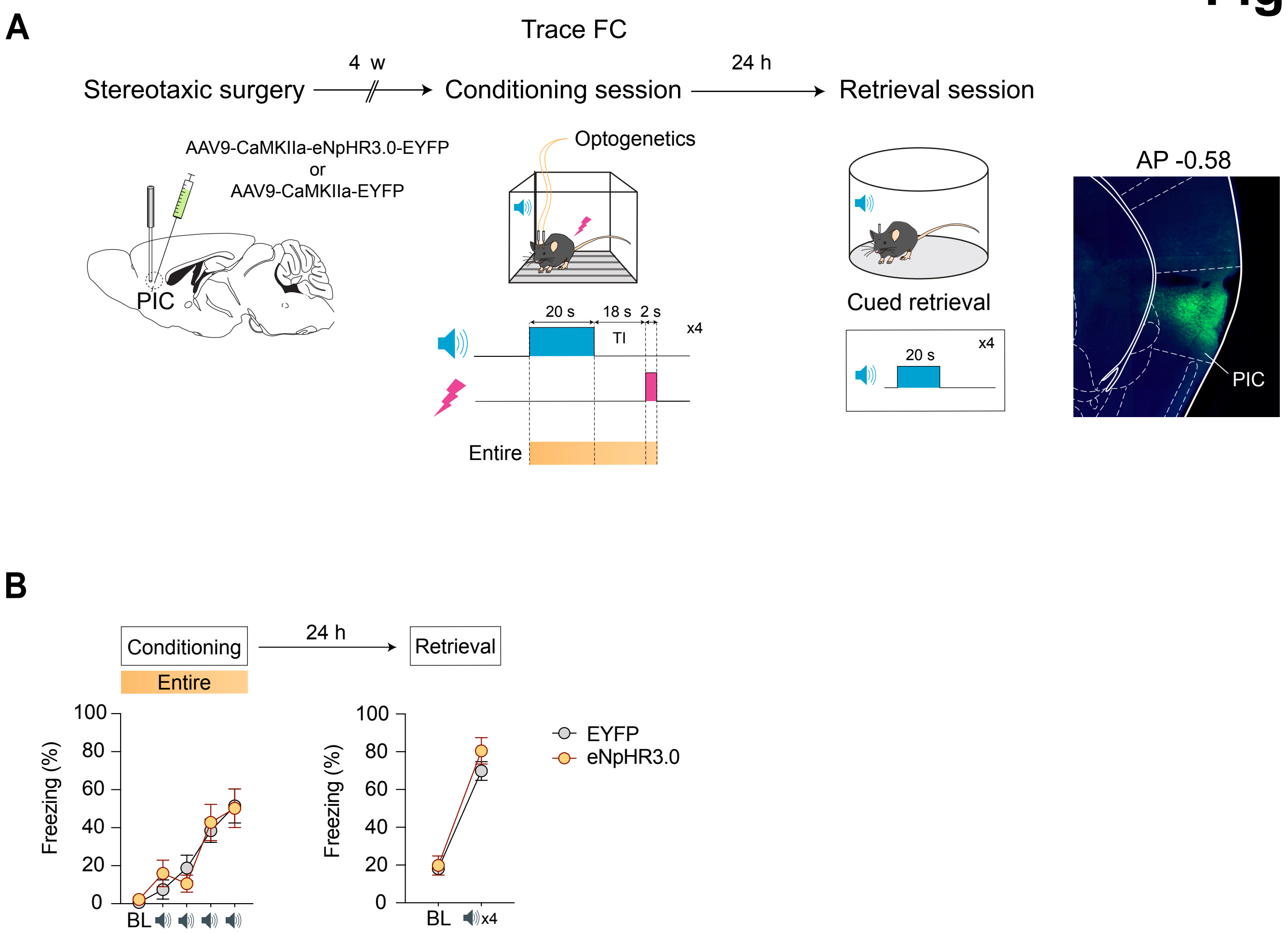
A
Trace FC
Stereotaxic surgery
4 w
Conditioning session
24 h
Retrieval session
AAV9-CaMKIIa-eNpHR3.0-EYFP
or
AAV9-CaMKIIa-EYFP
PIC
Optogenetics
20 s
18 s
2 s
TI
x4
Entire
Cued retrieval
20 s
x4
AP -0.58
PIC
B
Conditioning
24 h
Retrieval
Entire
Freezing (%)
100
80
60
40
20
0
BL
x4
EYFP
eNpHR3.0

**Figure S3. Activity of PIC excitatory neurons is not required for fear memory acquisition in trace FC.**

**(A)** Schematic of the optogenetic inhibition experiments targeting PIC excitatory neurons.

**(B)** Optogenetic inhibition of PIC excitatory neurons throughout the entire CS-TI-US period did not affect fear memory acquisition in trace FC. Conditioning ($N_{EYFP}$=10, $N_{eNpHR3.0}$=11; two-way mixed ANOVA, $p$=0.630). Retrieval ($N_{EYFP}$=10, $N_{eNpHR3.0}$=11; two-way mixed ANOVA, $p$=0.286).

More detailed statistical information is in Table S1.

### Fig. S4

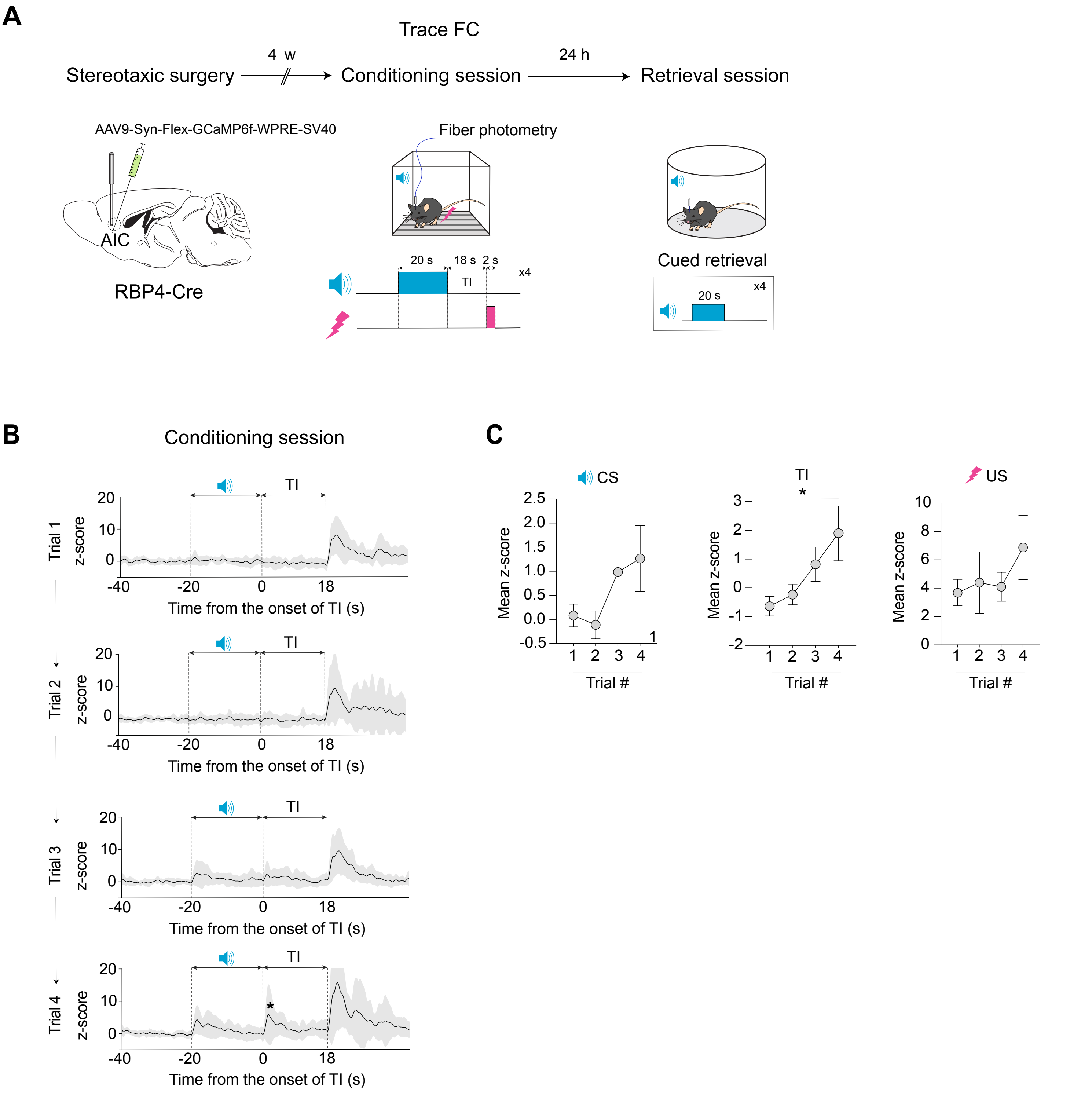
A
Trace FC
Stereotaxic surgery
4 w
Conditioning session
24 h
Retrieval session
AAV9-Syn-Flex-GCaMP6f-WPRE-SV40
AIC
RBP4-Cre
Fiber photometry
20 s
18 s
2 s
TI
x4
Cued retrieval
20 s
x4
B
Conditioning session
Trial 1
Trial 2
Trial 3
Trial 4
z-score
TI
Time from the onset of TI (s)
C
CS
TI
US
Mean z-score
Trial #

**Figure S4. AIC L5-PNs show a learning-related activity at the onset of the TI that becomes increasingly prominent across the trials during the conditioning session.**

**(A)** Schematic of the fiber photometry experiments recording population-level activity in AIC L5-PNs.

**(B)** Recordings revealed a transient activity (asterisk) time-locked to the onset of the TI across four CS-US pairing trials (Trial 1 - Trial4).

**(C)** During the conditioning session, mean activity significantly increased during the TI, but not during the CS or the US, across the trials. CS ($N$=10; one-way repeated measures ANOVA, $p$=0.092). TI ($N$=10; one-way repeated measures ANOVA, $p$=0.029). US ($N$=10; one-way repeated measures ANOVA, $p$=0.220). *$p < 0.05$, Data are expressed as mean ± SEM. More detailed statistical information is in Table S1.

### Fig. S5

**A**

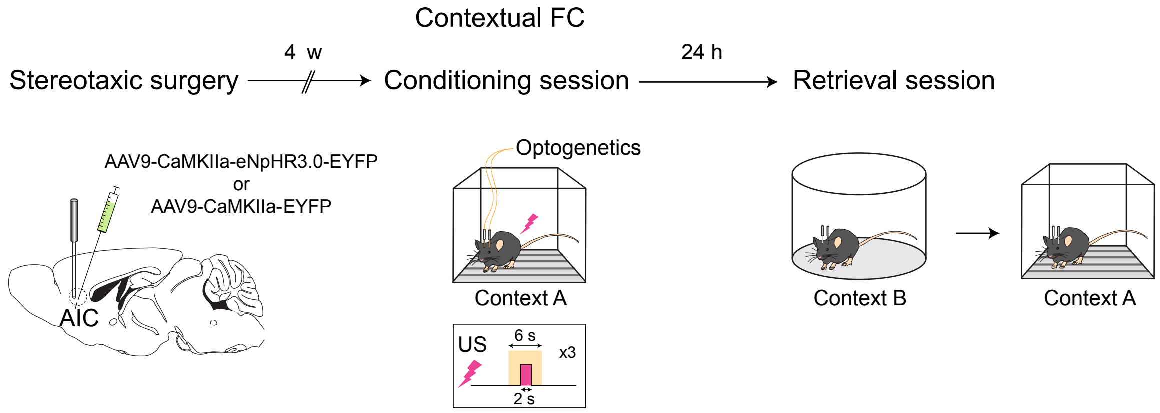
Contextual FC
Stereotaxic surgery
4 w
Conditioning session
24 h
Retrieval session
AAV9-CaMKIIa-eNpHR3.0-EYFP
or
AAV9-CaMKIIa-EYFP
AIC
Optogenetics
Context A
US
6 s
x3
2 s
Context B
Context A

**B**

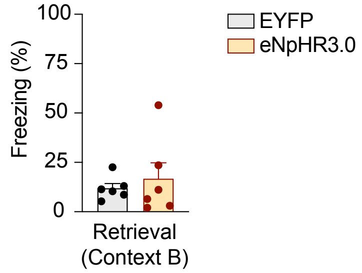
100
75
50
25
0
Freezing (%)
EYFP
eNpHR3.0
Retrieval
(Context B)

**Figure S5. Optogenetic inhibition of AIC excitatory neurons during the conditioning session does not affect fear responses to a neutral context.**

**(A)** Schematic of temporally specific optogenetic inhibition experiments during the conditioning session.

**(B)** Optogenetic inhibition of AIC excitatory neurons during the 6-second time windows that surround each 2-second US did not affect fear responses to a neutral context (Context B) during the retrieval session. Retrieval ($N_{EYFP}$=6, $N_{eNpHR3.0}$=6; two-tailed Student's *t*-test, $p$=0.582). More detailed statistical information is in Table S1.

### Fig. S6

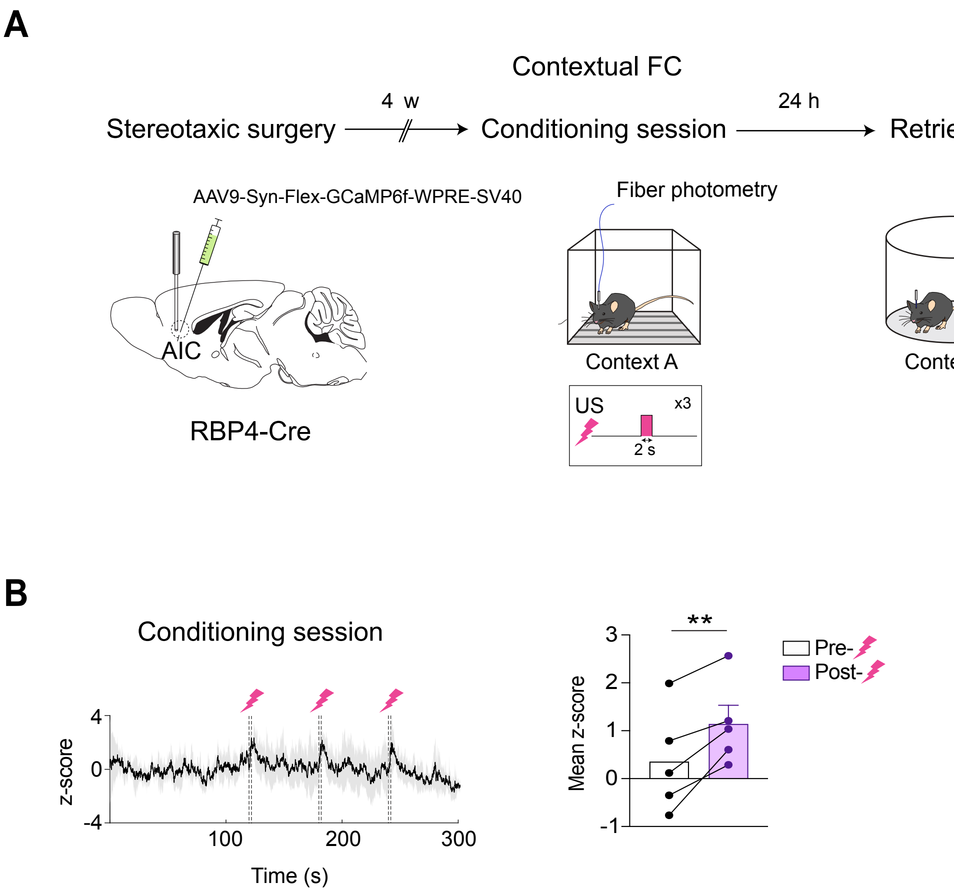
A
Contextual FC
Stereotaxic surgery
4 w
Conditioning session
24 h
Retrieval session
AAV9-Syn-Flex-GCaMP6f-WPRE-SV40
AIC
RBP4-Cre
Fiber photometry
Context A
US
x3
2 s
Context B
Context A
B
Conditioning session
z-score
4
0
-4
100
200
300
Time (s)
Mean z-score
3
2
1
0
-1
**
Pre-
Post-

**Figure S6. AIC L5-PNs exhibit strong responses to the US during the conditioning session in contextual FC.**

**(A)** Schematic of the fiber photometry experiments recording from AIC L5-PNs.

**(B)** Left: Population-level recordings revealed strong responses to US delivery across three context-US pairings. Right: Mean activity in the 10-s window following US onset was significantly higher than in the 10-s pre-US baseline window. mean z-score ($N = 5$; Two-tailed paired Student's $t$-test; $p = 0.009$). **$p < 0.01$. Data are expressed as mean ± SEM. More detailed statistical information is in Table S1.

### Fig. S7

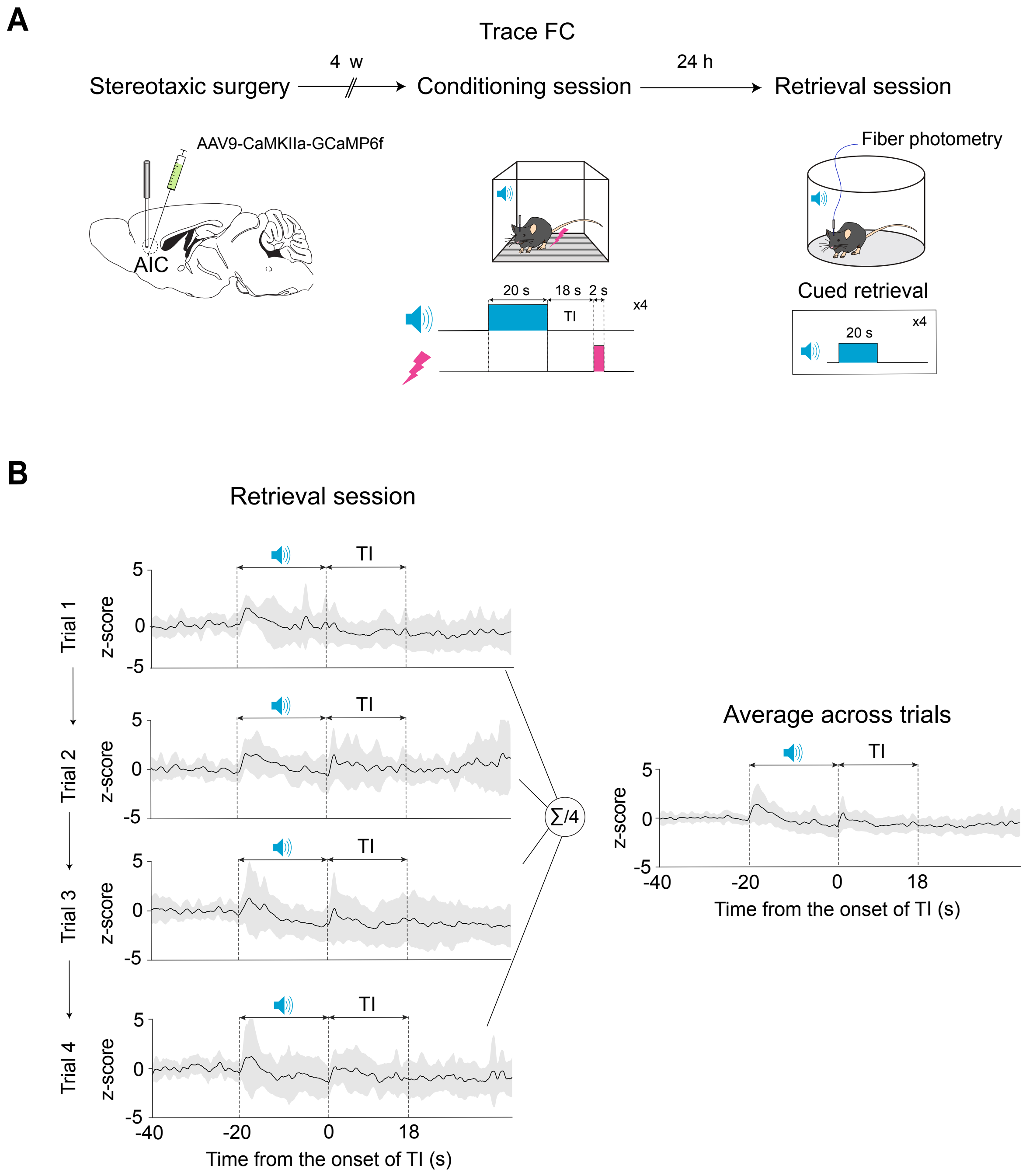
A
Trace FC
Stereotaxic surgery
4 w
Conditioning session
24 h
Retrieval session
AAV9-CaMKIIa-GCaMP6f
AIC
Fiber photometry
20 s
18 s
2 s
TI
x4
Cued retrieval
20 s
x4
B
Retrieval session
Trial 1
Trial 2
Trial 3
Trial 4
z-score
TI
5
0
-5
-40
-20
0
18
Time from the onset of TI (s)
∑/4
Average across trials
Time from the onset of TI (s)

**Figure S7. During the retrieval session in trace FC, AIC excitatory neurons maintain consistent responses to the CS and the TI across trials.**

**(A)** Schematic of the fiber photometry experiments recording from AIC excitatory neurons.

**(B)** Population-level recordings of AIC excitatory neurons showed their consistent responses to the CS and the TI during the retrieval session in trace FC.

### Fig. S8

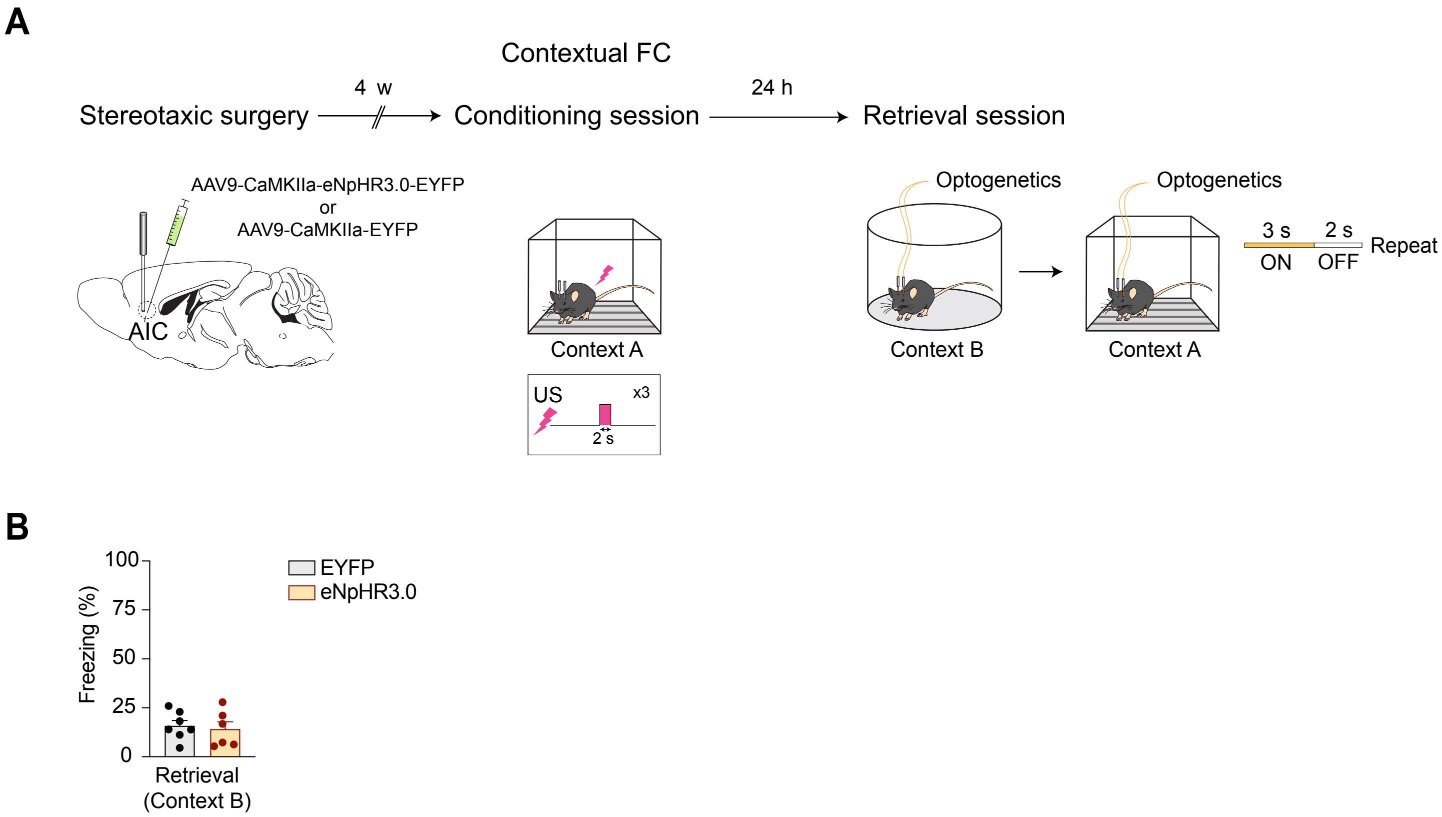
A
Contextual FC
Stereotaxic surgery
4 w
Conditioning session
24 h
Retrieval session
AAV9-CaMKIIa-eNpHR3.0-EYFP
or
AAV9-CaMKIIa-EYFP
AIC
Context A
US
x3
2 s
Optogenetics
Optogenetics
Context B
Context A
3 s
2 s
ON
OFF
Repeat
B
Freezing (%)
100
75
50
25
0
EYFP
eNpHR3.0
Retrieval
(Context B)

**Figure S8. Optogenetic inhibition of AIC excitatory neurons during the retrieval session does not affect fear responses to a neutral context.**

**(A)** Schematic of temporally specific optogenetic inhibition experiments during the retrieval session.

**(B)** Optogenetic inhibition of AIC excitatory neurons during the retrieval session on a 3-second ON/2-second OFF cycle did not alter fear responses to a neutral context (Context B). Retrieval ($N_{EYFP}$=7, $N_{eNpHR3.0}$=6; two-tailed Student's *t*-test, $p$=0.717). More detailed statistical information is in Table S1.

### Fig. S9

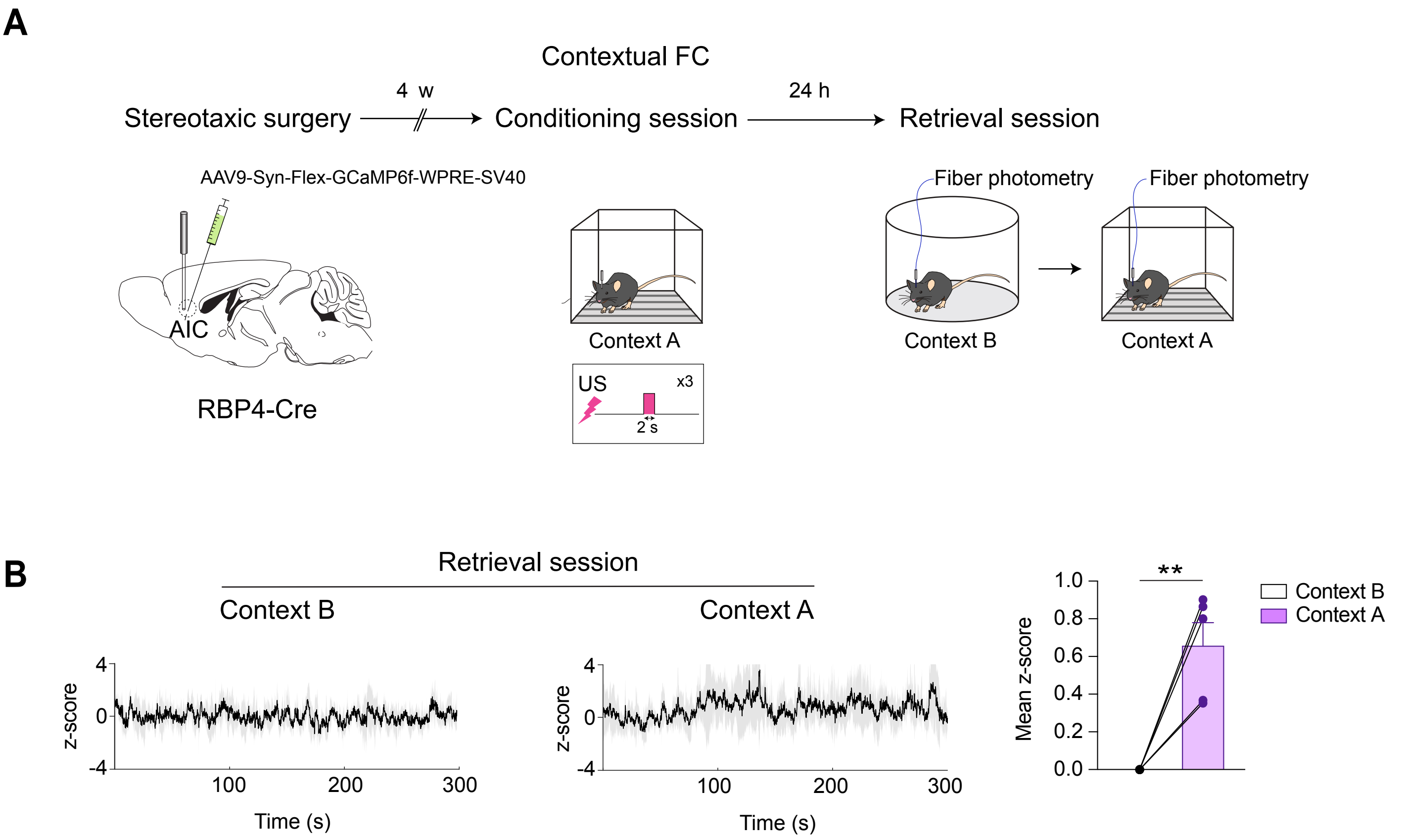

A
Contextual FC
Stereotaxic surgery
4 w
Conditioning session
24 h
Retrieval session
AAV9-Syn-Flex-GCaMP6f-WPRE-SV40
AIC
RBP4-Cre
Context A
US
x3
2 s
Fiber photometry
Context B
Fiber photometry
Context A
B
Retrieval session
Context B
Context A
z-score
4
0
-4
100
200
300
Time (s)
z-score
4
0
-4
100
200
300
Time (s)
**
Mean z-score
1.0
0.8
0.6
0.4
0.2
0.0
Context B
Context A

**Figure S9. AIC L5-PNs exhibit strong responses to the conditioning context during the retrieval session.**

**(A)** Schematic of the fiber photometry experiments recording from AIC L5-PNs.

**(B)** Left: During the retrieval session, AIC L5-PNs exhibit elevated, context-driven activity when mice were exposed to the conditioning context (Context A) but not to a neutral context (Context B). Right: Mean activity was significantly greater during exposure to the conditioning context compared to a neutral context. mean z-score ($N = 5$; Two-tailed one-sample $t$-test; $p = 0.006$). $^{**}p < 0.01$. Data are expressed as mean ± SEM. More detailed statistical information is in Table S1.

### Fig. S10

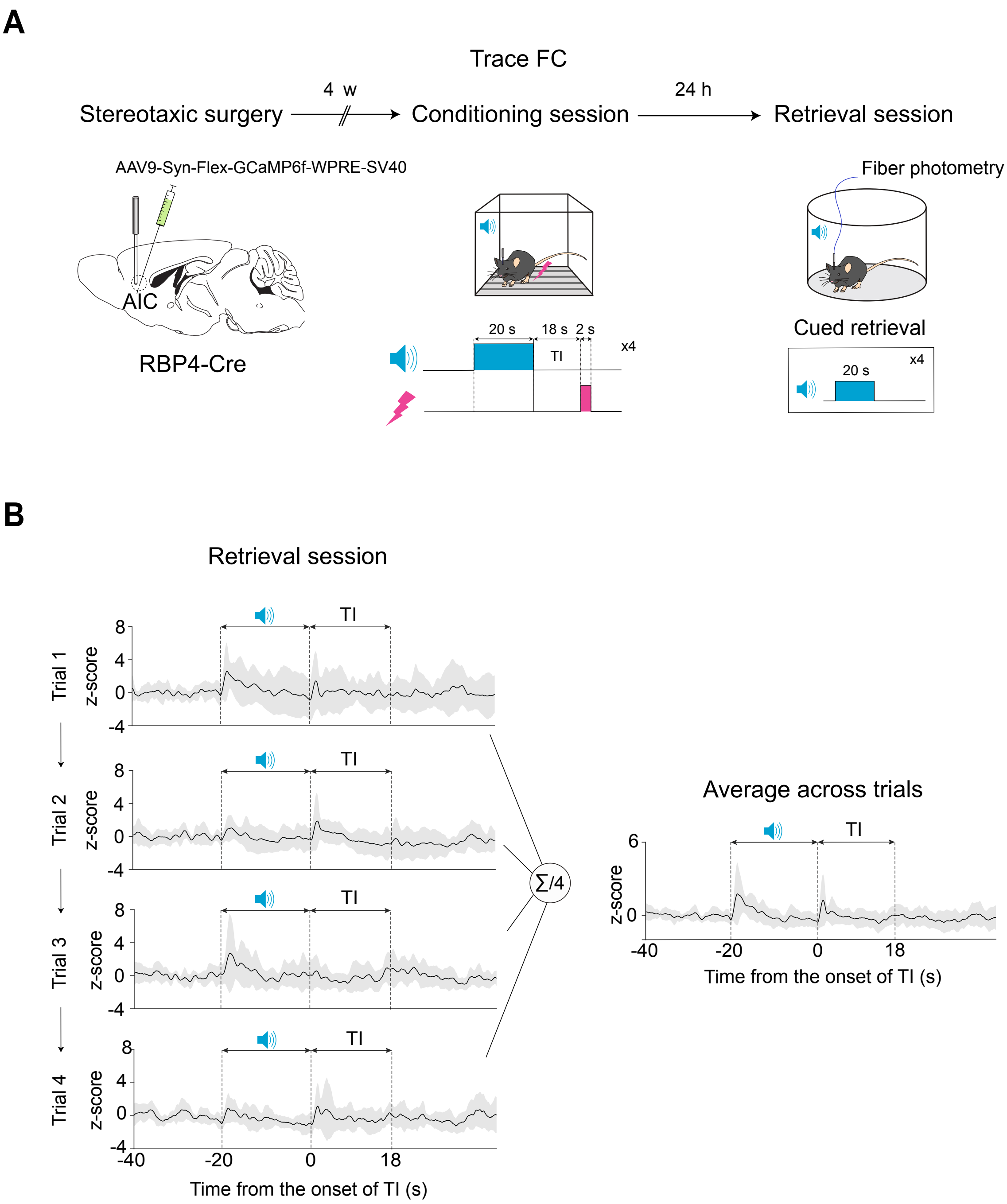
A
Trace FC
Stereotaxic surgery
4 w
Conditioning session
24 h
Retrieval session
AAV9-Syn-Flex-GCaMP6f-WPRE-SV40
AIC
RBP4-Cre
20 s
18 s
2 s
TI
x4
Fiber photometry
Cued retrieval
20 s
x4
B
Retrieval session
Trial 1
Trial 2
Trial 3
Trial 4
z-score
TI
∑/4
Average across trials
Time from the onset of TI (s)
-40
-20
0
18

**Figure S10. During the retrieval session in trace FC, AIC L5-PNs maintain consistent responses to the CS and the TI across trials.**

**(A)** Schematic of the fiber photometry experiments recording from AIC L5-PNs.

**(B)** Population-level recordings of AIC L5-PNs showed their consistent responses to the CS and the TI during the retrieval session in trace FC.

### Fig. S11

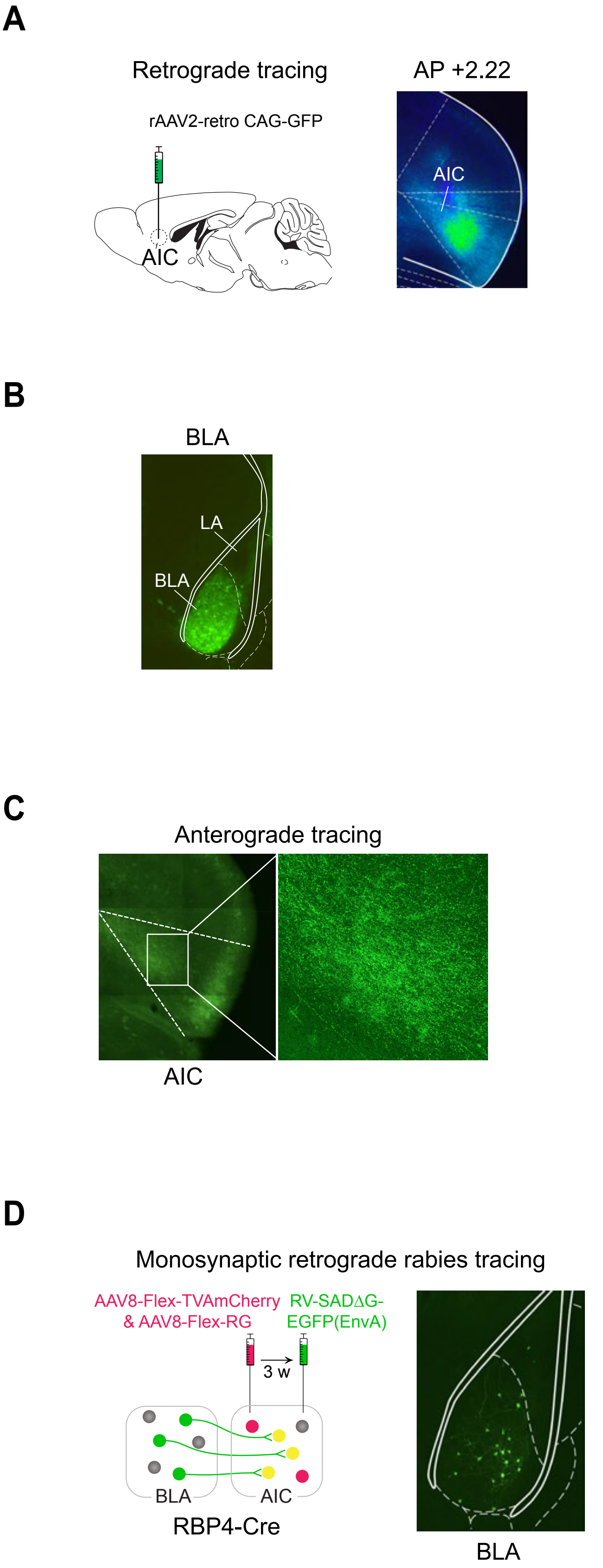
A
Retrograde tracing
rAAV2-retro CAG-GFP
AIC
AP +2.22
AIC
B
BLA
LA
BLA
C
Anterograde tracing
AIC
D
Monosynaptic retrograde rabies tracing
AAV8-Flex-TVAmCherry
& AAV8-Flex-RG
RV-SADΔG-
EGFP(EnvA)
3 w
BLA
AIC
RBP4-Cre
BLA

**Figure S11. Viral tracing reveals robust inputs from the BLA to AIC L5-PNs.**

**(A)** AAV-mediated retrograde tracing used to map afferent inputs to the AIC.

**(B)** Retrograde transport of GFP into the AIC revealed enriched populations of AIC-projecting neurons in the BLA.

**(C)** Anterograde labeling of BLA axons demonstrated dense terminal fields in layer 5.

**(D)** Cell type-specific, rabies virus-based retrograde tracing from AIC L5-PNs further confirmed BLA neurons that provide monosynaptic inputs.

**Table S1 Statistics**

| Measurement | Group | Sub-group | Sample size | Average | Standard deviation | Statistical test | Statistical result | Post hoc test | Post hoc test result |
|---|---|---|---|---|---|---|---|---|---|
| Freezing (%) | Vehicle | BL | 12 | 13.190 | 10.566 | Two-way Mixed ANOVA with the Greenhouse-Geisser correction<br>Between-subjects factor: Drug (vehicle vs. M+B)<br>Within-subjects factor: CS | Drug: $F$(1,22)=0.122, $p$ =0.731<br>CS: $F$(3.027,66.584)=58.086, $p$ =1.225E-18<br>Drug*CS: $F$(3.027,66.584)=1.455, $p$ =0.235 | - | - |
| | | CS1 | | 33.200 | 28.584 | | | | |
| | | CS2 | | 42.488 | 24.423 | | | | |
| | | CS3 | | 58.967 | 25.055 | | | | |
| | | CS4 | | 72.558 | 16.389 | | | | |
| | M+B | BL | 12 | 10.770 | 9.223 | | | | |
| | | CS1 | | 26.250 | 23.396 | | | | |
| | | CS2 | | 47.896 | 26.023 | | | | |
| | | CS3 | | 72.858 | 17.272 | | | | |
| | | CS4 | | 73.217 | 16.777 | | | | |
| Freezing (%) | Vehicle | BL | 12 | 13.737 | 8.125 | Two-way Mixed ANOVA<br>Between-subjects factor: Drug (vehicle vs. M+B)<br>Within-subjects factor: CS | Drug: $F$(1,22)=1.417, $p$ =0.247<br>CS: $F$(1,22)=57.243, $p$ =1.472E-07<br>Drug*CS: $F$(1,22)=0.703, $p$ =0.411 | - | - |
| | | CS (average) | | 58.275 | 20.939 | | | | |
| | M+B | BL | 12 | 12.667 | 7.564 | | | | |
| | | CS (average) | | 48.317 | 25.049 | | | | |
| Freezing (%) | Vehicle | BL | 11 | 8.787 | 14.416 | Two-way Mixed ANOVA with the Greenhouse-Geisser correction<br>Between-subjects factor: Drug (vehicle vs. M+B)<br>Within-subjects factor: CS | Drug: $F$(1,22)=0.075, $p$ =0.787<br>CS: $F$(2.946,64.802)=54.468, $p$ =1.491E-17<br>Drug*CS: $F$(2.946,64.802)=0.596, $p$ =0.617 | - | - |
| | | CS1 | | 22.927 | 19.309 | | | | |
| | | CS2 | | 47.109 | 28.108 | | | | |
| | | CS3 | | 72.591 | 29.007 | | | | |
| | | CS4 | | 90.059 | 19.252 | | | | |
| | M+B | BL | 13 | 12.579 | 17.373 | | | | |
| | | CS1 | | 26.915 | 30.502 | | | | |
| | | CS2 | | 44.404 | 40.935 | | | | |
| | | CS3 | | 61.065 | 36.351 | | | | |
| | | CS4 | | 85.177 | 21.415 | | | | |
| Freezing (%) | Vehicle | BL | 11 | 9.977 | 12.308 | Two-way Mixed ANOVA<br>Between-subjects factor: Drug (vehicle vs. M+B)<br>Within-subjects factor: CS | Drug: $F$(1,22)=3.276, $p$ =0.084<br>CS: $F$(1,22)=56.336, $p$ =1.674E-07<br>Drug*CS: $F$(1,22)=7.888, $p$ =0.010 | - | - |
| | | CS (average) | | 58.439 | 24.438 | | | | |
| | M+B | BL | 13 | 10.388 | 10.763 | | | | |
| | | CS (average) | | 32.458 | 28.853 | | | | |
| Freezing (%) | Vehicle | BL | 10 | 5.727 | 5.249 | Two-way Mixed ANOVA with the Greenhouse-Geisser correction<br>Between-subjects factor: Drug (vehicle vs. M+B)<br>Within-subjects factor: CS | Drug: $F$(1,20)=0.087, $p$ =0.771<br>CS: $F$(2.787,55.744)=55.414, $p$ =3.224E-16<br>Drug*CS: $F$(2.787,55.744)=1.084, $p$ =0.360 | - | - |
| | | CS1 | | 19.035 | 26.247 | | | | |
| | | CS2 | | 48.360 | 27.738 | | | | |
| | | CS3 | | 66.915 | 17.164 | | | | |
| | | CS4 | | 75.020 | 14.545 | | | | |
| | M+B | BL | 12 | 9.292 | 9.927 | | | | |
| | | CS1 | | 27.104 | 20.626 | | | | |
| | | CS2 | | 37.250 | 19.290 | | | | |
| | | CS3 | | 66.883 | 12.460 | | | | |
| | | CS4 | | 68.308 | 19.522 | | | | |
| Freezing (%) | Vehicle | BL | 10 | 12.360 | 11.738 | Two-way Mixed ANOVA<br>Between-subjects factor: Drug (vehicle vs. M+B)<br>Within-subjects factor: CS | Drug: $F$(1,20)=0.056, $p$ =0.816<br>CS: $F$(1,20)=55.105, $p$ =3.630E-07<br>Drug*CS: $F$(1,20)=1.646, $p$ =0.214 | - | - |
| | | CS (average) | | 63.105 | 24.303 | | | | |
| | M+B | BL | 12 | 18.020 | 19.495 | | | | |
| | | CS (average) | | 53.809 | 29.537 | | | | |
| Freezing (%) | Vehicle | BL | 10 | 7.456 | 2.781 | Two-way Mixed ANOVA with the Greenhouse-Geisser correction<br>Between-subjects factor: Drug (vehicle vs. M+B)<br>Within-subjects factor: CS | Drug: $F$(1,18)=0.010, $p$ =0.920<br>CS: $F$(2.105,37.899)=50.853, $p$ =1.079E-11<br>Drug*CS: $F$(2.105,37.899)=2.134, $p$ =0.130 | - | - |
| | | CS1 | | 27.483 | 7.778 | | | | |
| | | CS2 | | 33.268 | 10.277 | | | | |
| | | CS3 | | 59.033 | 21.634 | | | | |
| | | CS4 | | 62.599 | 31.177 | | | | |
| | M+B | BL | 10 | 3.981 | 1.685 | | | | |
| | | CS1 | | 14.067 | 6.642 | | | | |
| | | CS2 | | 30.583 | 10.777 | | | | |
| | | CS3 | | 59.465 | 31.350 | | | | |
| | | CS4 | | 79.701 | 18.816 | | | | |
| Freezing (%) | Vehicle | BL | 10 | 15.471 | 9.550 | Two-way Mixed ANOVA<br>Between-subjects factor: Drug (vehicle vs. M+B)<br>Within-subjects factor: CS | Drug: $F$(1,18)=0.566, $p$ =0.462<br>CS: $F$(1,18)=88.237, $p$ =2.319E-08<br>Drug*CS: $F$(1,18)=0.355, $p$ =0.559 | - | - |
| | | CS (average) | | 70.955 | 16.066 | | | | |
| | M+B | BL | 10 | 22.462 | 8.583 | | | | |
| | | CS (average) | | 71.325 | 25.933 | | | | |
| Freezing (%) | EYFP | BL | 10 | 14.0500 | 16.70304 | Two-way Mixed ANOVA with the Greenhouse-Geisser<br>Between-subjects factor: AAV (EYFP vs. eNpHR3.0)<br>Within-subjects factor: CS | AAV: $F$(1,18)=0.256, $p$ =0.619<br>CS: $F$(2.430,43.736)=13.537, $p$ =8.507E-06<br>AAV*CS: $F$(2.430,43.736)=0.589, $p$ =0.591 | - | - |
| | | CS1 | | 17.6300 | 29.74749 | | | | |
| | | CS2 | | 26.9300 | 30.96482 | | | | |
| | | CS3 | | 37.4400 | 30.57712 | | | | |
| | | CS4 | | 53.4200 | 32.19896 | | | | |
| | eNpHR3.0 | BL | 10 | 7.6900 | 8.37675 | | | | |
| | | CS1 | | 7.4900 | 7.77695 | | | | |
| | | CS2 | | 21.1100 | 25.24561 | | | | |
| | | CS3 | | 46.1800 | 26.80571 | | | | |
| | | CS4 | | 47.0600 | 34.13780 | | | | |
| Freezing (%) | EYFP | BL | 10 | 10.9500 | 5.39861 | Two-way Mixed ANOVA<br>Between-subjects factor: AAV (EYFP vs. eNpHR3.0)<br>Within-subjects factor: CS | AAV: $F$(1,18)=8.993, $p$ =0.008<br>CS: $F$(1,18)=86.416, $p$ =2.713E-08<br>AAV*CS: $F$(1,18)=8.225, $p$ =0.009 | - | - |
| | | CS (average) | | 66.4900 | 26.09523 | | | | |
| | eNpHR3.0 | BL | 10 | 7.5300 | 7.30389 | | | | |
| | | CS (average) | | 36.8800 | 16.12705 | | | | |

| Measurement | Group | Sub-group | Sample size | Average | Standard deviation | Statistical test | Statistical result | Post hoc test | Post hoc test result |
|---|---|---|---|---|---|---|---|---|---|
| Freezing (%) | EYFP | BL | 10 | 7.050 | 6.910 | Two-way Mixed ANOVA with the Greenhouse-Geisser correction<br>Between-subjects factor: AAV (EYFP vs. eNpHR3.0)<br>Within-subjects factor: CS | AAV: $F$ (1,18)=0.142, $p$ =0.711<br>CS: $F$ (2.572,46.294)=25.706, $p$ =2.890E-09<br>AAV*CS: $F$ (2.572,46.294)=0.201, $p$ =0.868 | - | - |
| | | CS1 | | 7.430 | 9.643 | | | | |
| | | CS2 | | 32.010 | 23.714 | | | | |
| | | CS3 | | 59.330 | 42.070 | | | | |
| | | CS4 | | 62.350 | 37.163 | | | | |
| | eNpHR3.0 | BL | 10 | 12.610 | 13.325 | | | | |
| | | CS1 | | 15.740 | 18.649 | | | | |
| | | CS2 | | 36.920 | 28.599 | | | | |
| | | CS3 | | 59.590 | 36.041 | | | | |
| | | CS4 | | 59.740 | 33.783 | | | | |
| Freezing (%) | EYFP | BL | 10 | 11.500 | 16.037 | Two-way Mixed ANOVA<br>Between-subjects factor: AAV (EYFP vs. eNpHR3.0)<br>Within-subjects factor: CS | AAV: $F$ (1,18)=5.081, $p$ =0.037<br>CS: $F$ (1,18)=36.388, $p$ =1.055E-05<br>AAV*CS: $F$ (1,18)=5.919, $p$ =0.026 | - | - |
| | | CS (average) | | 66.720 | 27.294 | | | | |
| | eNpHR3.0 | BL | 10 | 9.850 | 8.699 | | | | |
| | | CS (average) | | 33.330 | 31.329 | | | | |
| Freezing (%) | EYFP | BL | 11 | 13.782 | 14.590 | Two-way Mixed ANOVA<br>Between-subjects factor: AAV (EYFP vs. eNpHR3.0)<br>Within-subjects factor: CS | AAV: $F$ (1,20)=0.651, $p$ =0.429<br>CS: $F$ (4,80)=30.510, $p$ =2.026E-15<br>AAV*CS: $F$ (4,80)=0.550, $p$ =0.700 | - | - |
| | | CS1 | | 27.109 | 27.425 | | | | |
| | | CS2 | | 48.391 | 26.769 | | | | |
| | | CS3 | | 56.336 | 32.940 | | | | |
| | | CS4 | | 73.055 | 27.080 | | | | |
| | eNpHR3.0 | BL | 11 | 9.782 | 8.516 | | | | |
| | | CS1 | | 14.845 | 15.555 | | | | |
| | | CS2 | | 34.564 | 31.169 | | | | |
| | | CS3 | | 57.982 | 28.957 | | | | |
| | | CS4 | | 71.027 | 33.013 | | | | |
| Freezing (%) | EYFP | BL | 11 | 21.955 | 23.534 | Two-way Mixed ANOVA<br>Between-subjects factor: AAV (EYFP vs. eNpHR3.0)<br>Within-subjects factor: CS | AAV: $F$ (1,20)=0.239, $p$ =0.630<br>CS: $F$ (1,20)=76.578, $p$ =2.837E-08<br>AAV*CS: $F$ (1,20)=0.170, $p$ =0.685 | - | - |
| | | CS (average) | | 70.282 | 19.835 | | | | |
| | eNpHR3.0 | BL | 11 | 20.627 | 19.884 | | | | |
| | | CS (average) | | 64.609 | 19.935 | | | | |
| Freezing (%) | EYFP | BL | 11 | 19.073 | 21.776 | Two-way Mixed ANOVA<br>Between-subjects factor: AAV (EYFP vs. eNpHR3.0)<br>Within-subjects factor: CS | AAV: $F$ (1,20)=0.360, $p$ =0.555<br>CS: $F$ (4,80)=34.810, $p$ =8.099E-17<br>AAV*CS: $F$ (4,80)=0.897, $p$ =0.470 | - | - |
| | | CS1 | | 29.127 | 19.513 | | | | |
| | | CS2 | | 39.864 | 22.453 | | | | |
| | | CS3 | | 65.855 | 23.745 | | | | |
| | | CS4 | | 70.564 | 19.673 | | | | |
| | eNpHR3.0 | BL | 11 | 15.764 | 11.474 | | | | |
| | | CS1 | | 22.591 | 20.658 | | | | |
| | | CS2 | | 46.427 | 26.049 | | | | |
| | | CS3 | | 56.245 | 34.965 | | | | |
| | | CS4 | | 60.609 | 25.921 | | | | |
| Freezing (%) | EYFP | BL | 11 | 18.009 | 19.678 | Two-way Mixed ANOVA<br>Between-subjects factor: AAV (EYFP vs. eNpHR3.0)<br>Within-subjects factor: CS | AAV: $F$ (1,20)=0.943, $p$ =0.343<br>CS: $F$ (1,20)=48.837, $p$ =8.807E-07<br>AAV*CS: $F$ (1,20)=2.470, $p$ =0.132 | - | - |
| | | CS (average) | | 66.755 | 25.434 | | | | |
| | eNpHR3.0 | BL | 11 | 19.036 | 17.202 | | | | |
| | | CS (average) | | 49.882 | 29.107 | | | | |
| Mean z-score | CS | CS1 | 12 | -0.090 | 1.018 | One-way Repeated Measures ANOVA<br>Repeated factor: CS | CS: $F$ (3,33)=0.731, $p$ =0.552 | | |
| | | CS2 | | -0.263 | 1.476 | | | | |
| | | CS3 | | 0.125 | 0.688 | | | | |
| | | CS4 | | 0.347 | 1.037 | | | | |
| | TI | TI1 | | -0.852 | 1.366 | One-way Repeated Measures ANOVA<br>Repeated factor: TI | TI: $F$ (3,33)=3.996, $p$ =0.016 | | |
| | | TI2 | | -0.660 | 1.068 | | | | |
| | | TI3 | | -0.032 | 0.615 | | | | |
| | | TI4 | | 0.549 | 0.997 | | | | |
| | US | US1 | | 3.468 | 3.391 | One-way Repeated Measures ANOVA<br>Repeated factor: US | US: $F$ (3,33)=1.732, $p$ =0.179 | | |
| | | US2 | | 2.132 | 2.635 | | | | |
| | | US3 | | 4.629 | 3.114 | | | | |
| | | US4 | | 3.705 | 2.742 | | | | |
| GFP intensity (a.u.) | Layer | L1 | 3 | 1.946 | 0.448 | One-way ANOVA | $F$ (3,8)=33.346, $p$ =7.176E-05 | Tukey's HSD | $p$ (L1 vs. L2/3)=0.073,<br>$p$ (L1 vs. L5)=7.498E-05<br>$p$ (L1 vs. L6)=0.661<br>$p$ (L2/3 vs. L5)=0.001<br>$p$ (L2/3 vs. L6)=0.356<br>$p$ (L5 vs. L6)=2.011E-04 |
| | | L2/3 | | 9.990 | 3.662 | | | | |
| | | L5 | | 27.119 | 5.296 | | | | |
| | | L6 | | 5.156 | 1.886 | | | | |

| Measurement | Group | Sub-group | Sample size | Average | Standard deviation | Statistical test | Statistical result | Post hoc test | Post hoc test result |
|---|---|---|---|---|---|---|---|---|---|
| Freezing (%) | mCherry | BL | 8 | 9.200 | 12.456 | Two-way Mixed ANOVA with the Greenhouse-Geisser correction<br>Between-subjects factor: AAV (mCherry vs. hM4D(Gi))<br>Within-subjects factor: CS | AAV: *F* (1,14)=0.464, *p* =0.507<br>CS: *F* (2.720,38.084)=25.702, *p* =6.810E-09<br>AAV*CS: *F* (2.720,38.084)=1.550, *p* =0.220 | - | - |
| | | CS1 | | 21.675 | 20.876 | | | | |
| | | CS2 | | 35.113 | 24.283 | | | | |
| | | CS3 | | 72.413 | 32.788 | | | | |
| | | CS4 | | 59.675 | 34.681 | | | | |
| | hM4D(Gi) | BL | 8 | 15.913 | 14.867 | | | | |
| | | CS1 | | 21.275 | 19.999 | | | | |
| | | CS2 | | 60.913 | 25.932 | | | | |
| | | CS3 | | 66.250 | 25.906 | | | | |
| | | CS4 | | 62.913 | 23.557 | | | | |
| Freezing (%) | mCherry | BL | 8 | 14.125 | 9.097 | Two-way Mixed ANOVA<br>Between-subjects factor: AAV (mCherry vs. hM4D(Gi))<br>Within-subjects factor: CS | AAV: *F* (1,14)=2.522, *p* =0.135<br>CS: *F* (1,14)=93.559, *p* =1.409E-07<br>AAV*CS: *F* (1,14)=5.336, *p* =0.037 | - | - |
| | | CS (average) | | 56.338 | 9.315 | | | | |
| | hM4D(Gi) | BL | 8 | 13.688 | 17.162 | | | | |
| | | CS (average) | | 39.625 | 14.176 | | | | |
| Freezing (%) | EYFP | BL | 10 | 18.080 | 14.054 | Two-way Mixed ANOVA<br>Between-subjects factor: AAV (EYFP vs. eOPN3)<br>Within-subjects factor: CS | AAV: *F* (1,18)=0.004, *p* =0.951<br>CS: *F* (4,72)=18.956, *p* =1.102E-10<br>AAV*CS: *F* (4,72)=0.544, *p* =0.704 | - | - |
| | | CS1 | | 21.620 | 19.113 | | | | |
| | | CS2 | | 40.470 | 29.681 | | | | |
| | | CS3 | | 60.160 | 27.996 | | | | |
| | | CS4 | | 62.740 | 28.363 | | | | |
| | eOPN3 | BL | 10 | 13.160 | 12.052 | | | | |
| | | CS1 | | 25.880 | 33.876 | | | | |
| | | CS2 | | 48.700 | 33.470 | | | | |
| | | CS3 | | 51.690 | 33.981 | | | | |
| | | CS4 | | 60.690 | 35.168 | | | | |
| Freezing (%) | EYFP | BL | 10 | 15.570 | 16.852 | Two-way Mixed ANOVA<br>Between-subjects factor: AAV (EYFP vs. eOPN3)<br>Within-subjects factor: CS | AAV: *F* (1,18)=5.209, *p* =0.035<br>CS: *F* (1,18)=31.116, *p* =2.7041E-05<br>AAV*CS: *F* (1,18)=5.961, *p* =0.025 | - | - |
| | | CS (average) | | 59.240 | 17.797 | | | | |
| | eOPN3 | BL | 10 | 14.070 | 13.418 | | | | |
| | | CS (average) | | 31.150 | 25.546 | | | | |
| Freezing (%) | EYFP | BL | 12 | 4.500 | 6.029 | Two-way Mixed ANOVA<br>Between-subjects factor: AAV (EYFP vs. eOPN3)<br>Within-subjects factor: CS | AAV: *F* (1,22)=0.673, *p* =0.421<br>CS: *F* (2.373,52.213)=17.540, *p* =3.911E-07<br>AAV*CS: *F* (2.373,52.213)=0.093, *p* =0.937 | - | - |
| | | CS1 | | 8.650 | 13.862 | | | | |
| | | CS2 | | 22.167 | 24.842 | | | | |
| | | CS3 | | 38.725 | 32.025 | | | | |
| | | CS4 | | 47.942 | 37.081 | | | | |
| | eOPN3 | BL | 12 | 12.625 | 16.909 | | | | |
| | | CS1 | | 15.192 | 25.618 | | | | |
| | | CS2 | | 29.733 | 33.147 | | | | |
| | | CS3 | | 44.758 | 23.982 | | | | |
| | | CS4 | | 49.525 | 27.517 | | | | |
| Freezing (%) | EYFP | BL | 12 | 19.658 | 16.984 | Two-way Mixed ANOVA<br>Between-subjects factor: AAV (EYFP vs. eOPN3)<br>Within-subjects factor: CS | AAV: *F* (1,22)=0.421, *p* =0.523<br>CS: *F* (1,22)=61.694, *p* =7.994E-08<br>AAV*CS: *F* (1,22)=0.577, *p* =0.456 | - | - |
| | | CS (average) | | 65.975 | 28.690 | | | | |
| | eOPN3 | BL | 12 | 18.792 | 14.839 | | | | |
| | | CS (average) | | 56.942 | 27.575 | | | | |
| Freezing (%) | Vehicle | Before shock | 7 | 2.5129 | 2.15366 | Two-way Mixed ANOVA<br>Between-subjects factor: Drug (vehicle vs. M+B)<br>Within-subjects factor: Shock | Drug: *F* (1,12)=0.838, *p* =0.378<br>Shock: *F* (1,12)=45.575, *p* =2.041E-05<br>Drug*Shock: *F* (1,12)=0.600, | - | - |
| | | After shock | | 33.5414 | 19.01110 | | | | |
| | M+B | Before shock | 7 | 2.6900 | 4.70005 | | | | |
| | | After shock | | 41.7586 | 16.83481 | | | | |
| Freezing (%) | Vehicle | - | 7 | 44.1571 | 14.07520 | Two-tailed Student's *t* -test | *t* (12)=4.245, *p* =0.001 | - | - |
| | M+B | - | 7 | 15.9214 | 10.56161 | | | | |
| Freezing (%) | Vehicle | Before shock | 7 | 7.009 | 4.827 | Two-way Mixed ANOVA<br>Between-subjects factor: Drug (vehicle vs. M+B)<br>Within-subjects factor: Shock | Drug: *F* (1,12)=0.030, *p* =0.866<br>Shock: *F* (1,12)=43.969, *p* =2.426E-05<br>Drug*Shock: *F* (1,12)=0.079, | - | - |
| | | After shock | | 33.526 | 11.509 | | | | |
| | M+B | Before shock | 7 | 4.880 | 3.421 | | | | |
| | | After shock | | 33.737 | 22.512 | | | | |
| Freezing (%) | Vehicle | - | 7 | 46.373 | 19.125 | Two-tailed Student's *t* -test | *t* (12)=-3.667, *p* =0.003 | - | - |
| | M+B | - | 7 | 82.891 | 18.127 | | | | |
| Mean z-score | Shock | No-shock | 6 | -0.087 | 0.874 | Two-tailed paired Student's t-test | *t* (5)=-7.377, *p* =7.196E-04 | - | - |
| | | Shock | | 2.301 | 1.194 | | | | |
| Freezing (%) | EYFP | Before shock | 6 | 4.872 | 2.783 | Two-way Mixed ANOVA<br>Between-subjects factor: AAV (EYFP vs. eNpHR3.0)<br>Within-subjects factor: Shock | AAV: *F* (1,10)=0.010, *p* =0.924<br>Shock: *F* (1,10)=23.834, *p* =0.001<br>AAV*Shock: *F* (1,10)=0.058, *p* =0.814 | - | - |
| | | After shock | | 32.037 | 13.760 | | | | |
| | eNpHR3.0 | Before shock | 6 | 5.650 | 4.683 | | | | |
| | | After shock | | 30.260 | 20.685 | | | | |
| Freezing (%) | EYFP | - | 6 | 42.273 | 22.483 | Two-tailed Welch's *t* -test | *t* (5.656)=2.505, *p* =0.049 | - | - |
| | eNpHR3.0 | - | 6 | 18.537 | 5.769 | | | | |
| Freezing (%) | mCherry | Before shock | 5 | 7.780 | 10.878 | Two-way Mixed ANOVA<br>Between-subjects factor: AAV (mCherry vs. hM4D(Gi))<br>Within-subjects factor: Shock | AAV: *F* (1,9)=0.007, *p* =0.936<br>Shock: *F* (1,9)=9.319, *p* =0.014<br>AAV*Shock: *F* (1,9)=0.239, *p* =0.636 | - | - |
| | | After shock | | 20.060 | 15.197 | | | | |
| | hM4D(Gi) | Before shock | 6 | 4.900 | 4.813 | | | | |
| | | After shock | | 21.867 | 18.478 | | | | |
| Freezing (%) | mCherry | - | 5 | 59.780 | 20.693 | Two-tailed Student's t-test | *t* (9)=2.592, *p* =0.029 | - | - |
| | hM4D(Gi) | - | 6 | 27.500 | 20.471 | | | | |
| Freezing (%) | EYFP | Before shock | 6 | 4.472 | 4.446 | Two-way Mixed ANOVA<br>Between-subjects factor: AAV (EYFP vs. eOPN3)<br>Within-subjects factor: Shock | AAV: *F* (1,10)=0.016 *p* =0.903<br>Shock: *F* (1,10)=43.765, *p* =5.961E-05<br>AAV*Shock: *F* (1,10)=0.146, | - | - |
| | | After shock | | 42.415 | 17.771 | | | | |
| | eOPN3 | Before shock | 6 | 5.690 | 8.965 | | | | |
| | | After shock | | 39.495 | 22.216 | | | | |
| Freezing (%) | EYFP | - | 6 | 51.828 | 20.529 | Two-tailed Student's t-test | *t* (10)=2.382, *p* =0.0385 | - | - |
| | eOPN3 | - | 6 | 26.650 | 15.779 | | | | |

| Measurement | Group | Sub-group | Sample size | Average | Standard deviation | Statistical test | Statistical result | Post hoc test | Post hoc test result |
|---|---|---|---|---|---|---|---|---|---|
| Mean z-score | Context | Context B | 6 | 0.000 | 0.000 | Two-tailed one-sample t-test | *t* (5)=6.727, *p* =1.100E-03 | - | - |
| | | Context A | | 0.585 | 0.213 | | | | |
| Freezing (%) | EYFP | Before shock | 7 | 5.257 | 2.801 | Two-way Mixed ANOVA<br>Between-subjects factor: AAV (EYFP vs. eNpHR3.0)<br>Within-subjects factor: Shock | AAV: *F* (1,11)=0.195, *p* =0.667<br>Shock: *F* (1,11)=72.240, *p* =3.656E-06<br>AAV*Shock: *F* (1,11)=0.545, | - | - |
| | | After shock | | 40.886 | 14.267 | | | | |
| | eNpHR3.0 | Before shock | 6 | 5.667 | 5.322 | | | | - |
| | | After shock | | 35.600 | 19.042 | | | | |
| Freezing (%) | EYFP | - | 7 | 43.400 | 12.555 | Two-tailed Student's *t* -test | *t* (11)=-2.563, *p* =0.026 | - | - |
| | eNpHR3.0 | - | 6 | 66.650 | 19.895 | | | | |
| Freezing (%) | mCherry | Before shock | 5 | 3.750 | 4.024 | Two-way Mixed ANOVA<br>Between-subjects factor: AAV (mCherry vs. hM4D(Gi))<br>Within-subjects factor: Shock | AAV: *F* (1,8)=0.456, *p* =0.519<br>Shock: *F* (1,8)=29.971, *p* =0.001<br>AAV*Shock: *F* (1,8)=0.536, *p* =0.485 | - | - |
| | | After shock | | 31.690 | 18.943 | | | | |
| | hM4D(Gi) | Before shock | 5 | 10.994 | 6.217 | | | | |
| | | After shock | | 32.344 | 11.453 | | | | |
| Freezing (%) | mCherry | - | 5 | 57.048 | 13.858 | Two-tailed Student's t-test | *t* (8)=-2.355, *p* =0.046 | - | - |
| | hM4D(Gi) | - | 5 | 77.282 | 13.304 | | | | |
| Freezing (%) | EYFP | Before shock | 6 | 3.967 | 1.208 | Two-way Mixed ANOVA<br>Between-subjects factor: AAV (EYFP vs. eOPN3)<br>Within-subjects factor: Shock | AAV: *F* (1,10)=1.137 *p* =0.311<br>Shock: *F* (1,10)=104.703, *p* =1.287E-06<br>AAV*Shock: *F* (1,10)=0.174, | - | - |
| | | After shock | | 44.983 | 16.618 | | | | |
| | eOPN3 | Before shock | 6 | 10.600 | 7.333 | | | | |
| | | After shock | | 48.400 | 10.637 | | | | |
| Freezing (%) | EYFP | - | 6 | 52.883 | 16.623 | Two-tailed Student's t-test | *t* (10)=-2.399, *p* =0.037 | - | - |
| | eOPN3 | - | 6 | 75.600 | 16.173 | | | | |
| Cell proportion (%) | | Conditioning-only | 4 | 0.339 | 0.066 | One-way ANOVA | *F* (2,9)=38.649, *p* =3.820×10$^{-5}$ | Tukey's HSD | *p* (Conditioning-only vs. overlap)=0.003, *p* (Conditioning-only vs. Retrieval-only)=0.006 *p* (overlap vs. Retrieval-only)=2.762E-05 |
| | | Overlap | | 0.081 | 0.060 | | | | |
| | | Retrieval-only | | 0.580 | 0.109 | | | | |
| Freezing (%) | EYFP | Before shock | 5 | 16.498 | 12.833 | Two-way Mixed ANOVA<br>Between-subjects factor: AAV (EYFP vs. NpHR)<br>Within-subjects factor: Shock | AAV: *F* (1,9)=0.375, *p* =0.555<br>Shock: *F* (1,9)=63.265, *p* =2.318E-05<br>AAV*Shock: *F* (1,9)=0.008, *p* =0.931 | - | - |
| | | After shock | | 51.016 | 11.676 | | | | |
| | NpHR | Before shock | 6 | 13.573 | 12.818 | | | | |
| | | After shock | | 47.327 | 7.858 | | | | |
| Freezing (%) | EYFP | - | 5 | 46.884 | 9.108 | Two-tailed Student's t-test | *t* (9)=-1.624, *p* =0.139 | - | - |
| | NpHR | - | 6 | 55.295 | 8.081 | | | | |
| Freezing (%) | Vehicle | BL | 9 | 6.818 | 7.328 | Two-way Mixed ANOVA with the Greenhouse-Geisser correction<br>Between-subjects factor: Drug (vehicle vs. M+B)<br>Within-subjects factor: CS | Drug: *F* (1,16)=0.605, *p* =0.448<br>CS: *F* (2.701,43.215)=61.384, *p* =4.988E-15<br>Drug*CS: *F* (2.701,43.215)=0.138, *p* =0.922 | - | - |
| | | CS1 | | 11.427 | 16.603 | | | | |
| | | CS2 | | 22.630 | 28.276 | | | | |
| | | CS3 | | 57.445 | 31.985 | | | | |
| | | CS4 | | 73.536 | 21.407 | | | | |
| | M+B | BL | 9 | 9.577 | 11.263 | | | | |
| | | CS1 | | 13.908 | 13.613 | | | | |
| | | CS2 | | 30.296 | 23.107 | | | | |
| | | CS3 | | 65.352 | 29.803 | | | | |
| | | CS4 | | 81.740 | 19.372 | | | | |
| Freezing (%) | Vehicle | BL | 9 | 17.351 | 22.680 | Two-way Mixed ANOVA<br>Between-subjects factor: Drug (vehicle vs. M+B)<br>Within-subjects factor: CS | Drug: *F* (1,16)=0.042, *p* =0.840<br>CS: *F* (1,16)=35.862, *p* =1.893E-05<br>Drug*CS: *F* (1,16)=0.323, *p* =0.587 | - | - |
| | | CS (average) | | 49.890 | 28.935 | | | | |
| | M+B | BL | 9 | 15.732 | 16.860 | | | | |
| | | CS (average) | | 55.093 | 19.651 | | | | |
| Freezing (%) | mCherry | BL | 9 | 2.124 | 3.217 | Two-way Mixed ANOVA with the Greenhouse-Geisser correction<br>Between-subjects factor: AAV (mCherry vs. hM4D(Gi))<br>Within-subjects factor: CS | AAV: *F* (1,16)=2.446, *p* =0.137<br>CS: *F* (2.030,32.481)=103.014, *p* =7.381E-15<br>AAV*CS: *F* (2.030,32.481)=1.161, *p* =0.326 | - | - |
| | | CS1 | | 1.223 | 2.433 | | | | |
| | | CS2 | | 21.168 | 22.745 | | | | |
| | | CS3 | | 56.466 | 23.160 | | | | |
| | | CS4 | | 68.148 | 12.239 | | | | |
| | hM4D(Gi) | BL | 9 | 1.961 | 3.445 | | | | |
| | | CS1 | | 3.946 | 6.412 | | | | |
| | | CS2 | | 27.837 | 28.815 | | | | |
| | | CS3 | | 73.762 | 25.607 | | | | |
| | | CS4 | | 81.724 | 9.058 | | | | |
| Freezing (%) | mCherry | BL | 9 | 20.109 | 13.760 | Two-way Mixed ANOVA<br>Between-subjects factor: AAV (mCherry vs. hM4D(Gi))<br>Within-subjects factor: CS | AAV: *F* (1,16)=2.482, *p* =0.135<br>CS: *F* (1,16)=84.577, *p* =8.694E-08<br>AAV*CS: *F* (1,16)=0.625, *p* =0.441 | - | - |
| | | CS (average) | | 61.460 | 16.433 | | | | |
| | hM4D(Gi) | BL | 9 | 24.994 | 14.037 | | | | |
| | | CS (average) | | 74.121 | 18.369 | | | | |
| Freezing (%) | mCherry | Before shock | 8 | 5.936 | 2.146 | Two-way Mixed ANOVA<br>Between-subjects factor: AAV (mCherry vs. hM4D(Gi))<br>Within-subjects factor: Shock | AAV: *F* (1,15)=0.119, *p* =0.735<br>Shock: *F* (1,15)=44.493, *p* =7.476E-06<br>AAV*Shock: *F* (1,15)=1.703, | - | - |
| | | After shock | | 26.809 | 14.174 | | | | |
| | hM4D(Gi) | Before shock | 9 | 2.250 | 1.271 | | | | |
| | | After shock | | 33.276 | 17.802 | | | | |
| Freezing (%) | mCherry | - | 8 | 42.743 | 14.882 | Two-tailed Student's t-test | *t* (15)=-1.493, *p* =0.156 | - | - |
| | hM4D(Gi) | - | 9 | 55.068 | 18.644 | | | | |

| Measurement | Group | Sub-group | Sample size | Average | Standard deviation | Statistical test | Statistical result | Post hoc test | Post hoc test result |
|---|---|---|---|---|---|---|---|---|---|
| Freezing (%) | EYFP | BL | 10 | 0.660 | 0.992 | Two-way Mixed ANOVA<br>Between-subjects factor: AAV (EYFP vs. eNpHR3.0)<br>Within-subjects factor: CS | AAV: $F(1,19)=0.024$, $p=0.880$<br>CS: $F(4,76)=28.512$, $p=1.781\text{E-}14$<br>AAV*CS: $F(4,76)=0.648$, $p=0.630$ | - | - |
| | | CS1 | | 7.480 | 15.881 | | | | |
| | | CS2 | | 18.790 | 21.362 | | | | |
| | | CS3 | | 38.400 | 19.203 | | | | |
| | | CS4 | | 51.440 | 28.215 | | | | |
| | eNpHR3.0 | BL | 11 | 2.218 | 3.893 | | | | |
| | | CS1 | | 16.000 | 23.163 | | | | |
| | | CS2 | | 10.564 | 14.692 | | | | |
| | | CS3 | | 42.791 | 31.489 | | | | |
| | | CS4 | | 50.236 | 33.674 | | | | |
| Freezing (%) | EYFP | BL | 10 | 18.090 | 9.099 | Two-way Mixed ANOVA<br>Between-subjects factor: AAV (EYFP vs. eNpHR3.0)<br>Within-subjects factor: CS | AAV: $F(1,19)=0.972$, $p=0.337$<br>CS: $F(1,19)=192.685$, $p=2.135\text{E-}11$<br>AAV*CS: $F(1,19)=1.205$, $p=0.286$ | - | - |
| | | CS (average) | | 69.840 | 15.625 | | | | |
| | eNpHR3.0 | BL | 11 | 19.818 | 16.727 | | | | |
| | | CS (average) | | 80.455 | 23.076 | | | | |
| Mean z-score | CS | CS1 | 10 | 0.083 | 0.745 | One-way Repeated Measures ANOVA<br>Repeated factor: CS | CS: $F(3,27)=2.374$, $p=0.092$ | - | - |
| | | CS2 | | -0.113 | 0.914 | | | | |
| | | CS3 | | 0.984 | 1.642 | | | | |
| | | CS4 | | 1.267 | 2.154 | | | | |
| | TI | TI1 | | -0.632 | 1.084 | One-way Repeated Measures ANOVA<br>Repeated factor: TI | TI: $F(3,27)=3.505$, $p=0.029$ | - | - |
| | | TI2 | | -0.232 | 1.099 | | | | |
| | | TI3 | | 0.824 | 1.883 | | | | |
| | | TI4 | | 1.904 | 2.978 | | | | |
| | US | US1 | | 3.681 | 2.897 | One-way Repeated Measures ANOVA<br>Repeated factor: US | US: $F(3,27)=1.569$, $p=0.220$ | - | - |
| | | US2 | | 4.397 | 6.857 | | | | |
| | | US3 | | 4.104 | 3.230 | | | | |
| | | US4 | | 6.868 | 7.159 | | | | |
| Freezing (%) | EYFP | - | 6 | 11.877 | 5.876 | Two-tailed Student's $t$-test | $t(10)=-0.569$, $p=0.582$ | - | - |
| | eNpHR3.0 | - | 6 | 16.683 | 19.847 | | | | |
| Mean z-score | Shock | No-shock | 5 | 0.3564 | 1.08057 | Two-tailed paired Student's t-test | $t(4)=-4.685$, $p=0.009$ | - | - |
| | | Shock | | 1.1405 | 0.87346 | | | | |
| Freezing (%) | EYFP | - | 7 | 15.823 | 7.237 | Two-tailed Student's $t$-test | $t(11)=0.372$, $p=0.717$ | - | - |
| | eNpHR3.0 | - | 6 | 14.120 | 9.271 | | | | |
| Mean z-score | Context | Context B | 5 | 0.0000 | 0.00000 | Two-tailed one-sample t-test | $t(4)=5.364$, $p=0.006$ | - | - |
| | | Context A | | 0.6573 | 0.27399 | | | | |
